# A Spinal Circuit That Transmits Innocuous Cool Sensations

**DOI:** 10.1101/2022.03.23.485555

**Authors:** Lorraine R Horwitz, Hankyu Lee, Chia Chun Hor, Fred Y Shen, Wei Cai, Chen Li, Emily Ling-Lin Pai, Tin Long Rex Fung, Ilma Rovcanin, Kevin P Pipe, X.Z. Shawn Xu, Dawen Cai, Bo Duan

**Author notes:** Correspondence to: Bo Duan. These authors contributed equally (Lorraine R Horwitz, Hankyu Lee).

## Abstract

Temperature information is precisely processed in the nervous system. While progress has been made in identifying molecular thermosensors in the periphery, the neural circuits that process temperature information in the central nervous system remain unknown. Here we have identified an essential node in the neural circuitry for innocuous cool sensations. We found that a population of excitatory interneurons co-expressing Calbindin1 and Lbx1 (Calb1^Lbx1^) in the dorsal horn of the spinal cord is activated by innocuous cool temperatures. Genetic ablation or silencing of spinal Calb1^Lbx1^ neurons causes loss of innocuous cool but not noxious cold sensations. Further Brainbow labeling with expansion microscopy and electrophysiology showed that a small cluster of spinal Calb1^Lbx1^ interneurons in lamina I and the outer layer of lamina II represents the cooling-transmission neurons. These neurons receive monosynaptic connections from TRPM8^+^ primary sensory neurons and amplify the activity of cool-sensitive spinoparabrachial projection neurons. Our findings reveal a microcircuit in the dorsal spinal cord that specifically transmits innocuous cool sensations.

## INTRODUCTION

External temperatures are first detected in the skin and then transmitted via primary sensory neurons in the dorsal root ganglia (DRG) to the dorsal horn of the spinal cord for somatosensory integration (Montell and Caterina, 2007; Palkar et al., 2015; Patapoutian et al., 2003; Xiao and Xu, 2021). Thermal sensors for temperatures ranging from noxious heat/innocuous warm to innocuous cool/noxious cold allow the body to maintain homeostasis and protect against injury. Several members of the transient receptor potential (TRP) ion channel family have been classified as thermal sensors in mammals based on their ability to respond to a variety of stimuli ranging from noxious heat (TRPV1, TRPA1 and TRPM3) (Caterina et al., 2000; Caterina et al., 1997; Vandewauw et al., 2018) to innocuous cool (TRPM8) (Bautista et al., 2007; Colburn et al., 2007; Dhaka et al., 2007; McKemy et al., 2002; Milenkovic et al., 2014; Peier et al., 2002).

TRPM8 is a ligand-gated, nonselective cation channel that is activated by cool temperatures (22-27 °C) and by exogenous chemical agonists menthol and icilin (McKemy et al., 2002; Peier et al., 2002). TRPM8 channels are expressed in a subset of primary sensory neurons in the DRG (McKemy et al., 2002; Peier et al., 2002). Mice lacking *Trpm8* have severe deficits in the perception of innocuous cooling, but not noxious cold, suggesting the selective function of TRPM8 channels in the DRG for transmitting cool temperatures (Bautista et al., 2007; Colburn et al., 2007; Dhaka et al., 2007). TRPM8^+^ peripheral afferents project predominantly to laminae I and II outer layer (II_o_) of the spinal cord (Dhaka et al., 2008). However, the precise identity of the neural circuitry in the spinal cord that transmits cool information remains unknown.

The dorsal horn of the spinal cord is an integral component of the central nervous system for sensory processing. It consists of a complex network of local interneurons and projection neurons (Todd, 2010). *In vivo* single unit recordings and *in vivo* two-photon Ca2^+^ imaging studies have shown that a large population of interneurons and projection neurons in the superficial dorsal horn are thermosensitive (Andrew and Craig, 2001; Bester et al., 2000; Burton, 1975; Chisholm et al., 2021; Christensen and Perl, 1970; Craig and Kniffki, 1985; Craig et al., 2001; Ran et al., 2016). However, the lack of genetic tools and unique promoters to functionally manipulate specific neuronal subtypes makes it challenging to determine the cellular identities of thermosensitive neurons and circuits in the spinal cord.

In this study, using intersectional genetic manipulations (Duan et al., 2014; Pan et al., 2019), we have identified a population of excitatory interneurons co-expressing Calbindin1 (Calbindin-D28K, Calb1) and Lbx1 (hereafter referred to as Calb1^Lbx1^) in the dorsal horn that is essential to transmit innocuous cool but not noxious cold temperature information. Specifically, we found that a small cluster of Calb1^Lbx1^ interneurons in lamina I and II_o_, which does not express the neuropeptide somatostatin (SOM^-^), is required to transmit innocuous cool sensations. Alternatively, the subset of Calb1^Lbx1^ interneurons expressing SOM (SOM^+^) is required to transmit acute punctate mechanical pain. Finally, we showed that cool-sensitive TRPM8^+^ sensory neurons monosynapticially innervate Calb1^Lbx1^ interneurons in the superficial dorsal horn and that activation of Calb1^Lbx1^ interneurons amplifies the activity of cool-sensitive spinoparabrachial projection neurons (hereafter referred as SPB neurons). Taken together, these results delineate a spinal circuit that selectively transmits innocuous cool information to the brain.

## RESULTS

### Characterization of Calb1^Lbx1^ interneurons in the dorsal horn of the spinal cord

The calcium-binding protein Calb1 is expressed across the central nervous system including the DRG, trigeminal ganglion, amygdala, cerebellum, hippocampus, and brainstem (Sequier et al., 1990; Zhang et al., 1990). In the dorsal horn of the spinal cord, Calb1 is present in laminae I-IV (Zhang et al., 1990), suggesting Calb1 may act as a heterogenous marker for multiple sensations. Crossing *Calb1^Cre^* and *Lbx1^Flpo^* with a *Cre-* and *Flpo*-dependent Tomato reporter stain *Rosa26^CAG-ds-tdTomato^* (Ai65) (hereafter referred to as *Calb1^Lbx1^*;Ai65), enabled the selective expression of Tomato protein in Calb1^Lbx1^ neurons in the dorsal spinal cord.

Immunohistochemical double staining showed that 44.0% (961/2183) of Tomato^+^ neurons exhibited detectable Calb1 protein expression, and 85.3% (961/1126) of Calb1^Lbx1^ neurons co-expressed Tomato (Figure 1A).

**Figure 1.**
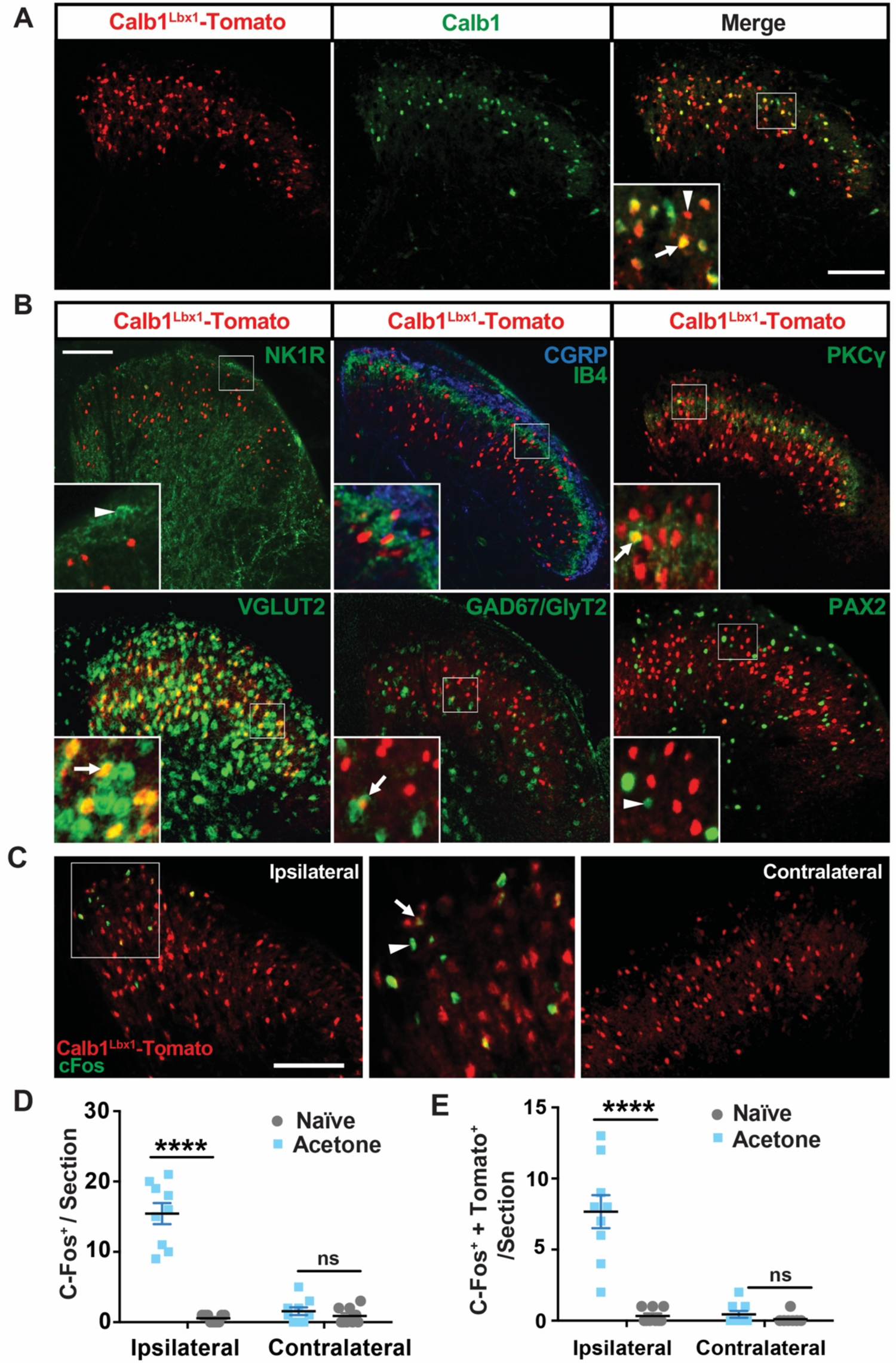
An innocuous Cool Stimulus Activates Calb1^Lbx1^-Tomato^+^ neurons in the Superficial Dorsal Horn of the Spinal Cord. **(A)** Images represent double staining of Calb1^Lbx1^-Tomato (red) and Calb1 protein (green) in *Calb1^Lbx1^*;Ai65 mice. Arrow denotes Calb1 protein and Tomato double-positive cell. Arrowhead indicates a Calb1^Lbx1^-Tomato positive cell that does not colocalize with Calb1 protein. n = 17 sections. Scale bar, 100 μm. **(B)** Double staining of Tomato with lamina markers (NK1R, CGRP, IB4, and PKCγ), excitatory neuronal marker VGLUT2, or inhibitory neuronal markers (GAD67 plus GlyT2 or Pax2) by immunohistochemistry or *in situ* hybridization in the dorsal horn of *Calb1^Lbx1^*;Ai65 mice. Arrows denote double-positive cells for indicated mRNA and Tomato. Arrowhead indicates a protein^+^ cell that does not colocalize with Tomato. The percentage is calculated as double-positive neurons over total number of Calb1^Lbx1^-Tomato^+^ neurons. Insets represent higher magnification of the boxed areas. n = 17-26 sections. Scale bar, 100 μm. **(C)** Double staining of c-Fos and Tomato signals in acetone-treated *Calb1^Lbx1^*;Ai65 mice. Inset (middle) represents higher magnification of the boxed area (left). Arrow indicates a double-positive cell for c-Fos protein and Tomato, and arrowhead shows a cell positive for c-Fos alone. Scale bar, 100 μm. **(D)** Total number of c-Fos positive neurons per hemi-section in either the ipsilateral or the contralateral dorsal horn of naïve (grey) and acetone-treated (light blue) *Calb1^Lbx1^*;Ai65 mice. n = 9 hemi-sections in each group; **** p < 0.0001 two-way ANOVA with Sidak post hoc analysis. **(E)** Quantification of c-Fos colocalization between c-Fos^+^ and Calb1^Lbx1^-Tomato^+^ neurons per hemi-section in either the ipsilateral or contralateral dorsal horn of naïve (grey) and acetone-treated (light blue) *Calb1^Lbx1^*;Ai65 mice. n = 9 hemi-sections in each group; **** p < 0.0001; two-way ANOVA with Sidak post hoc analysis.

To better characterize the cellular identity of Calb1^Lbx1^ neurons in the dorsal horn, sections of the lumbar spinal cord of *Calb1^Lbx1^;*Ai65 mice were immunostained with a panel of specific antibodies against markers of lamina layers. As shown in Figure 1B, we found a small portion of Tomato^+^ neurons were located in the lamina I (10.5%, 38/363) and very few Calb1^Lbx1^ neurons colocalized with Neurokinin 1 receptor (NK1R, a marker of a large population of ascending projection neurons in lamina I) (1.5%, 6/392, Figure 1B and S1). We also observed that Calb1^Lbx1^ neurons were intermingled with both calcitonin gene-related peptide (CGRP) terminals in laminae I-II_o_ (15.9%, 325/2039) and isolectin B4 (IB4) terminals in the dorsal part of lamina II inner layer (_d_II_i_) (23.4%, 742/3171) (Figure 1B). The largest populations of Calb1^Lbx1^ neurons were located in the ventral part of lamina II_i_ (_v_II_i_) to the dorsal part of lamina III (_d_III) (31.2%,1015/3253), which is marked by the expression of protein kinase C*γ* (PKC*γ*), and the ventral to the PKC*γ* zone in lamina III-IV (33.0%, 941/2855) (Figure 1B). To further identify types of Calb1^Lbx1^ neurons, *in situ* hybridization and immunohistochemistry approaches were used.

Figure 1B shows that the vast majority of Calb1^Lbx1^ neurons express vesicular glutamate transporter 2 (VGLUT2) mRNA (91.8%, 1692/1844), while only a small population of Calb1^Lbx1^ neurons express markers of inhibitory interneuron such as glutamic acid decarboxylase isoform 67 (GAD67)/glycine transporter 2 (GlyT2) mRNAs (9.0%, 180/2005) or paired box gene 2 (Pax2) protein (4.9%, 283/5793). These results indicate that Calb1^Lbx1^ neurons mainly represent a heterogeneous population of excitatory interneurons in laminae I-IV of the spinal cord.

### Innocuous cool temperatures activate Calb1^Lbx1^ neurons in the superficial dorsal horn

It is well established that acetone evaporation mimics cool temperatures, but not noxious cold temperatures in the range of 15-20 °C (Bautista et al., 2007). Here, we explored whether Calb1^Lbx1^ neurons are cool-sensitive. To do this, acetone was topically administered to the right hindpaw of *Calb1^Lbx1^*;Ai65 mice and the expression of the nuclear immediate early gene c-Fos in the lumbar spinal cord was assessed. We found that acetone treatment induced c-Fos protein expression in Tomato-expressing Calb1^Lbx1^ neurons in the ipsilateral spinal cord compared to naïve control mice (Figure 1C and 1D), and about half (49.6%, 69/139) of the Fos^+^ neurons were Tomato^+^ (Figure 1C and 1E). Interestingly, most Fos^+^;Tomato^+^ neurons were located in lamina I-II, suggesting that innocuous cool temperatures activate Calb1^Lbx1^ neurons in the superficial dorsal horn.

### Ablation of Calb1^Lbx1^ neurons results in innocuous cool sensing deficits

To further assess the function of Calb1^Lbx1^ neurons in transmitting innocuous cool sensations, we genetically ablated Calb1^Lbx1^ neurons in the dorsal spinal cord using an intersectional genetic strategy (Figure S2A), and then performed a battery of somatosensory behavioral paradigms. The ablation of Calb1^Lbx1^ neurons was performed by injecting *Calb1^Lbx1^;Tau^ds-DTR^* mice (which also carried a *Cre*-dependent Tomato reporter allele) with DTX (hereafter referred to as Calb1^Abl^ mice), resulting in a 91% reduction of Tomato^+^ neurons in the dorsal spinal cord (Figure 2A). It is worth mentioning that Calb1^Cre^ neurons in the brain, including somatosensation-related brain regions (the parabrachial nucleus and somatosensory cortex), were not ablated in Calb1^Abl^ mice (Figure S2B).

**Figure 2.**
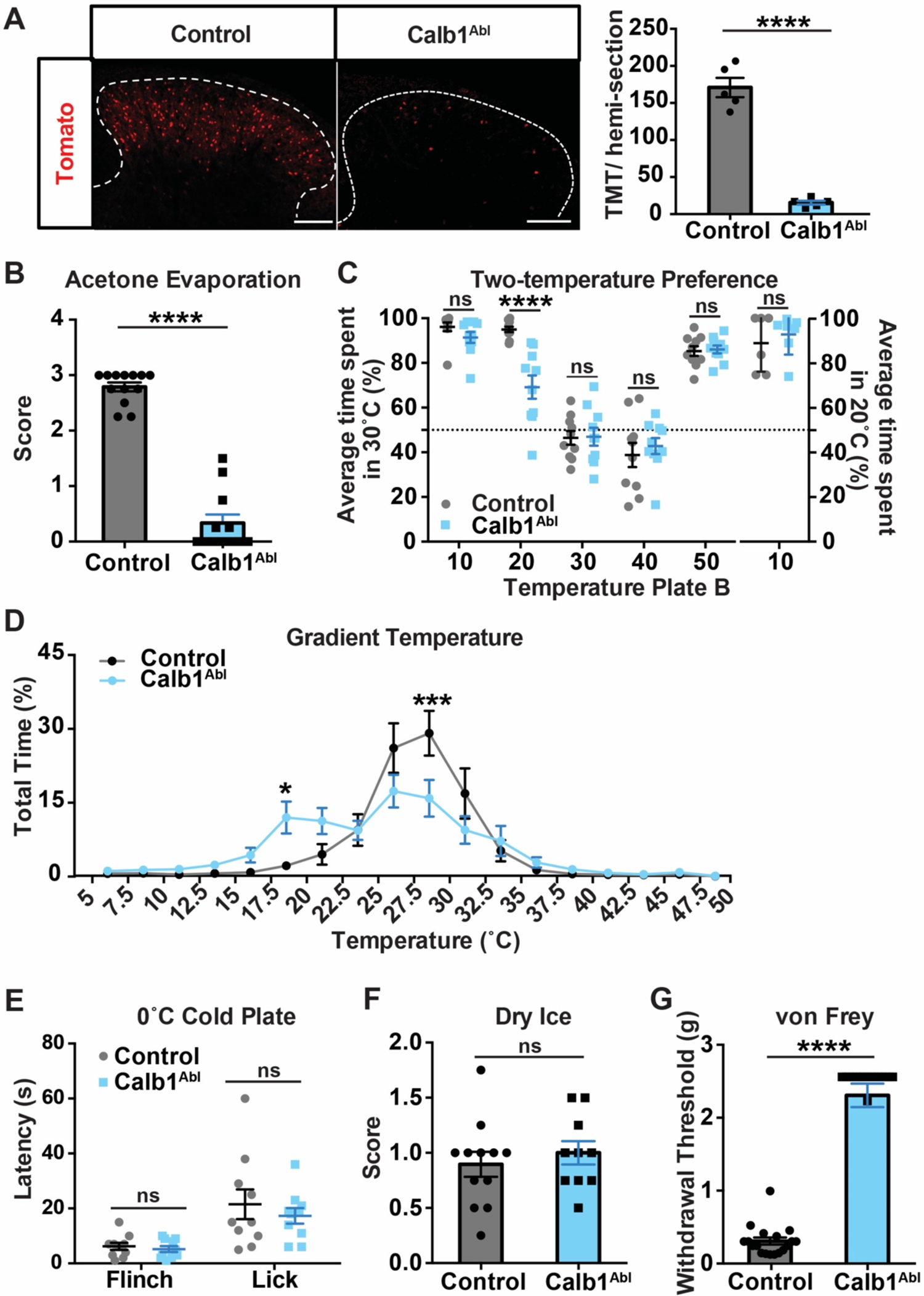
Loss of Innocuous Cool Sensations in Calb1^Abl^ Mice. **(A)** Ablation of Calb1^Lbx1^ neurons in the dorsal horn. Bar graph represents quantified data for Calb1^Cre^-Tomato signals in control and ablated animals. n = 27-30 sections; **** p < 0.0001, Student’s unpaired t test. Scale bar, 100 μm. **(B)** The acetone evaporation assay in Calb1^Abl^ and control groups. Control: n = 17; Calb1^Abl^: n = 14; **** p < 0.0001, Mann-Whitney test. **(C)** Two-temperature preference between two temperature plates. Left panel: Reference plate is set at 30 °C, and test plate temperature is set at 10 °C, 20 °C, 30 °C, 40 °C, and 50 °C. Control: n = 10-11; Calb1^Ab*l*^: n = 10; ns, no significant difference; **** p < 0.0001 two-way ANOVA with Sidak post hoc analysis. Right panel: Reference plate is set at 20 °C, test plate temperature is set at 10 °C. Data points represent the average percentage of time spent on the reference plate across two trials over the total trial time. Control: n = 6; Calb1^Abl^: n = 7; ns, no significant difference; Student’s unpaired t test. **(D)** Gradient temperature ranging from 5 °C to 50 °C is quantified as percentage of time spent in each temperature zone over the total trial time. Control: n = 13; Calb1^Abl^: n = 18; * p < 0.05; *** p < 0.001; two-way ANOVA with Bonferroni post hoc analysis. **(E)** Quantified forelimb withdrawal latency to 0 °C cold plate, including forelimb lick and flinch responses. n = 10 in each group; ns, no significant difference; Student’s unpaired t test. **(F)** Quantitative nocifensive response to dry ice application to the hindpaw. Score represents average response across four trials per mouse. Control: n = 12; Calb1^Abl^: n = 10; ns, no significant difference; Student’s unpaired t test. **(G)** Acute punctate mechanical pain threshold as measured by up-down von Frey withdrawal threshold was significantly increased in Calb1^Abl^ mice compared to controls. Control: n = 18; Calb1^Abl^: n = 13; **** p < 0.0001, Student’s unpaired t test.

To test locomotion in Calb1^Abl^ mice, we performed the rotarod assay and found that locomotor coordination remained intact in the ablated group (Figure S3A). Next, we assessed deficits in thermal temperature sensing in Calb1^Abl^ mice. As a first test, we asked whether acetone evaporation-induced cooling sensations were intact in Calb1^Abl^ mice. Following delivery of acetone to the plantar region of the hindpaw, we observed nocifensive responses that could be scored on a scale of 0 (no response or touch only without flinch) to 4 (noxious responses such as guarding, vocalizations, and escape behaviors). Compared to control littermates, Calb1^Abl^ mice showed nearly abolished responses to the innocuous cool stimuli (Figure 2B).

Next, we asked whether Calb1^Abl^ mice also exhibit deficits in their ability to discriminate between warm and cool using the two-temperature preference assay. Mice were given the choice between two temperature plates: a reference plate set to 30 °C and a test plate set to a fixed temperature from 10 °C to 50 °C. The percentage of time spent on the 30 °C surface was measured over a 5-minute period. When placed on equivalent temperatures (30 °C), both control and Calb1^Abl^ mice spent an equal amount of time on each plate (Figure 2C). When the test plate was set to 20 °C, control mice showed a clear preference for the reference plate at 30°C. By contrast, Calb1^Abl^ mice did not display a preference, spending nearly equal amounts of time on each side. Nonetheless, Calb1^Abl^ mice displayed a normal preference for 30 °C at noxious cold (10 °C), noxious heat (50 °C), and warm (40 °C) temperatures. Next, we compared the response of Calb1^Abl^ mice to 20 °C versus 10 °C, and found that Calb1^Abl^ mice did not show a deficit in this temperature-sensing test (Figure 2C).

To assess the thermal detection range of Calb1^Abl^ mice, the temperature gradient assay was performed. Mice were allowed to freely move across a surface temperature gradient of 5 °C to 50 °C. As shown in Figure 2D, Calb1^Abl^ mice spent significantly more time in the 17.5-20 °C temperature range and significantly less time in the 27.5-30 °C temperature range compared to controls, confirming innocuous cool sensing deficits. Neither control nor Calb1^Abl^ mice spent time in the temperature ranges of noxious cold (5-15 °C) and noxious heat (40-50 °C).

Consistently, when mice were exposed to a cold plate (0 °C), a hot plate (46, 50, or 54 °C), or received dry ice application to the hindpaw, there was no significant difference in nocifensive responses to noxious cold or noxious heat stimuli between Calb1^Abl^ and control mice (Figure 2E, 2F, and S3H). Furthermore, rectal temperature was not significantly different between control (37.0 ± 0.2 °C) and Calb1^Abl^ mice (36.9 ± 0.3 °C). Overall, temperature-sensing deficits were observed in Calb1^Abl^ mice exclusively at innocuous cool temperatures, suggesting Calb1^Lbx1^ neurons in the dorsal spinal cord play an essential role in innocuous cool sensing but not noxious cold.

Additional somatosensory assays were used to measure mechanosensitivity in control and Calb1^Abl^ mice. We found that ablation of Calb1^Lbx1^ neurons led to a dramatic increase in the threshold for acute punctate mechanical pain (Figure 2G and S3F). However, there was no significant difference in sharp mechanical pain as measured by the pinch and pinprick assays (Figure S3D and S3E). We also observed that Calb1^Abl^ mice displayed a decreased nocifensive response to the gentle touch of a paintbrush lightly brushed across the glabrous skin of the hindpaw (Figure S3B), but no difference in nocifensive responses to an innocuous sticky presentation to the glabrous skin of the hindpaw (Figure S3C).

### Silencing Calb1^Lbx1^ neurons results in innocuous cool deficits

To ensure that the observed behavioral results could not be attributed to potential secondary effects due to spinal circuits reorganization after neuronal ablation, we used intersectional strategies to transiently silence spinal Calb1^Lbx1^ neurons by crossing *Calb1^Cre^, Lbx1^Flpo^* with a *Cre-* and *Flpo*-dependent hM4Di designer receptors exclusively activated by a designer drug (DREADD) strain *Rosa26^CAG-ds-hM4Di^* (hereafter referred to as Calb1^Silenced^) (Figure S4A).

Clozapine N-oxide (CNO) was used to activate hM4Di thereby silencing spinal Calb1^Lbx1^ neurons, resulting in greatly attenuated responses to innocuous cool stimuli (Figure 3A-3C), whereas noxious cold sensations were unaffected (Figure 3D and 3E). Specifically, acute silencing of Calb1^Lbx1^ neurons resulted in a significant deficit in acetone-induced nocifensive responses (Figure 3A), compared to both baseline (before CNO treatment) and CNO-treated controls. In the two-temperature preference assay, Calb1^Silenced^ mice displayed a significant deficit in time spent in the 20 °C chamber, with no differences recorded at any other temperature (Figure 3B). Additionally, Calb1^Silenced^ mice spent significantly less time at the 27.5-30 °C temperature range, with the majority of time spent at cooler temperatures (Figure 3C).

**Figure 3.**
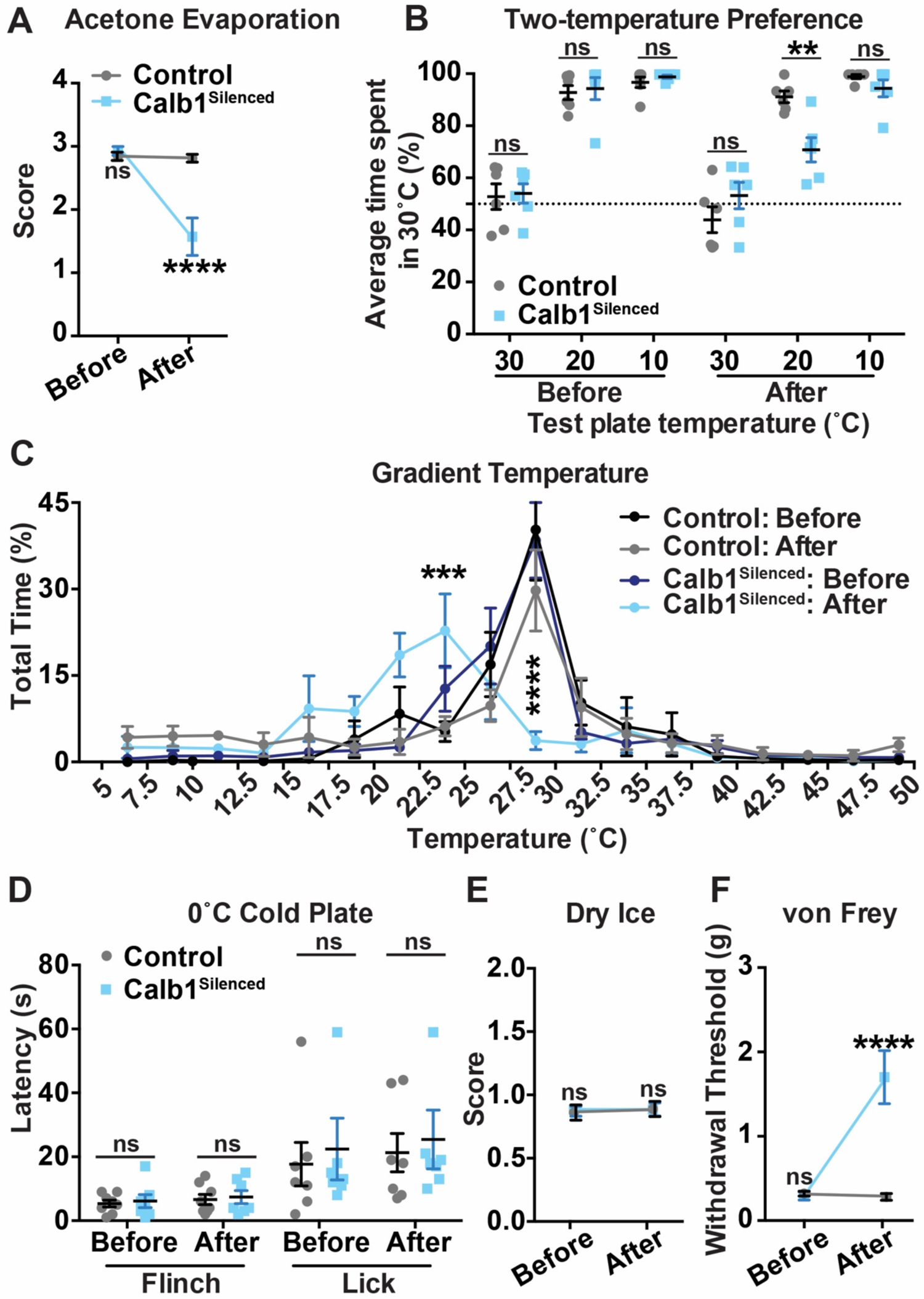
Silencing spinal Calb1^Lbx1^ Neurons Leads to Deficits in Innocuous Cool Sensations. **(A)** The acetone evaporation assay before and 40 minutes after CNO injection in mice with hM4Di receptors in spinal Calb1^Lbx1^-neuron silenced (Calb1^Silenced^) and control groups. Control: n = 8; Calb1^Silenced^: n = 7; **** p < 0.0001, two-way ANOVA with Sidak post hoc analysis. **(B)** Two-temperature preference between two temperature plates before and 40 minutes after CNO injection in Calb1^Silenced^ and control groups. Reference plate is set at 30°C, and test plate temperature is set at 30 °C, 20 °C and 10 °C. Data points represent the average percentage of time spent on the reference plate across two trials over the total trial time. n = 6 in each group; ** p < 0.01; two-way ANOVA with Sidak post hoc analysis. **(C)** Gradient temperature ranging from 5 °C to 50 °C was quantified as percentage of time spent in each temperature zone before and 40 minutes after CNO injection in Calb1^Silenced^ and control groups. Control: n = 4; Calb1^Silenced^: n = 7; *** p < 0.001; **** p < 0.0001; two-way ANOVA with Bonferroni post hoc analysis. **(D)** Quantified forelimb flinch and lick withdrawal latency to 0 °C cold plate before and 40 minutes after CNO injection in Calb1^Silenced^ and control groups. Control: n = 8; Calb1^Silenced^: n = 7; ns, no significant difference; two-way ANOVA with Sidak post hoc analysis. **(E)** Nocifensive responses to hindpaw application of dry ice stimulus before and 40 minutes after CNO injection in Calb1^Silenced^ and control groups. Score represents average response across four trials per mouse. Control: n = 9; Calb1^Silenced^: n = 11; two-way ANOVA with Sidak post hoc analysis. **(F)** Acute punctate mechanical pain measured using the up-down von Frey assay before and 40 minutes after CNO injection in Calb1^Silenced^ and control groups. Control: n = 8; Calb1^Silenced^: n = 7; **** p < 0.0001; two-way ANOVA with Sidak post hoc analysis.

While Calb1^Silenced^ mice displayed innocuous cool sensing behavior deficits, their ability to detect noxious cold stimuli was not affected in response to a 0 °C cold plate (Figure 3D), or dry ice application to the hindpaw (Figure 3E). Calb1^Sileneced^ mice also exhibited a significant increase in von Frey threshold (Figure 3F) and gentle touch deficits to light brushing (Figure S4C), while locomotion, innocuous touch, sharp mechanical pain, and thermal heat sensations remained intact (Figure S4B and S4D-S4H). Rectal temperature was not significantly different between control (37.3 ± 0.2 °C) and Calb1^Silenced^ (35.7 ± 1.2 °C) mice after CNO injection. Taken together, our results suggest that Calb1^Lbx1^ neurons represent a functionally diverse population in the dorsal spinal cord and are required to transmit innocuous cooling, acute punctate mechanical pain, and touch sensations.

### A subpopulation of Calb1^Lbx1^ neurons is responsible for innocuous cool sensing

To identify Calb1^Lbx1^ subpopulations responsible for innocuous cool sensing or acute punctate mechanical pain, we examined the overlap with a known marker of spinal excitatory interneurons, somatostatin (SOM) (Figure 4A). *In situ* hybridization experiments revealed that Calb1^Lbx1^ neurons highly overlapped with SOM (44%, 949/2157) in lamina II. Using the same intersectional genetic strategy, we ablated SOM^Lbx1^ neurons in the dorsal spinal cord and examined behavioral responses to thermal stimuli (Figure 4B-4D). We found that there was no significant difference between SOM^Abl^ and control mice in either the acetone evaporation assay (Figure 4B), the two-temperature preference assay (Figure 4C), or the temperature gradient assay (Figure 4D). Furthermore, noxious cold sensing remained intact (Figure 4C, 4D, and S5B). Consistently, few SOM^+^ neurons were activated after a cool stimulus was presented (Figure S5A). All together, these results suggest that SOM^+^ neurons are dispensable for cool sensing. Of note, SOM^Lbx1^ neurons are shown to be responsible for acute punctate mechanical pain (Duan et al., 2014), therefore the mechanical pain deficits exhibited in Calb1^Abl^ mice (Figure 2G), which are similar to the deficits in SOM^Abl^ mice (Figure 4E), could be attributed to Calb1^Lbx1^;SOM^+^ neurons. These results suggests that Calb1^Lbx1^;SOM^-^ neurons may form a subpopulation that transmits innocuous cool sensations, whereas Calb1^Lbx1^;SOM^+^ neurons may be required for sensing acute punctate mechanical pain (Figure 4F).

**Figure 4.**
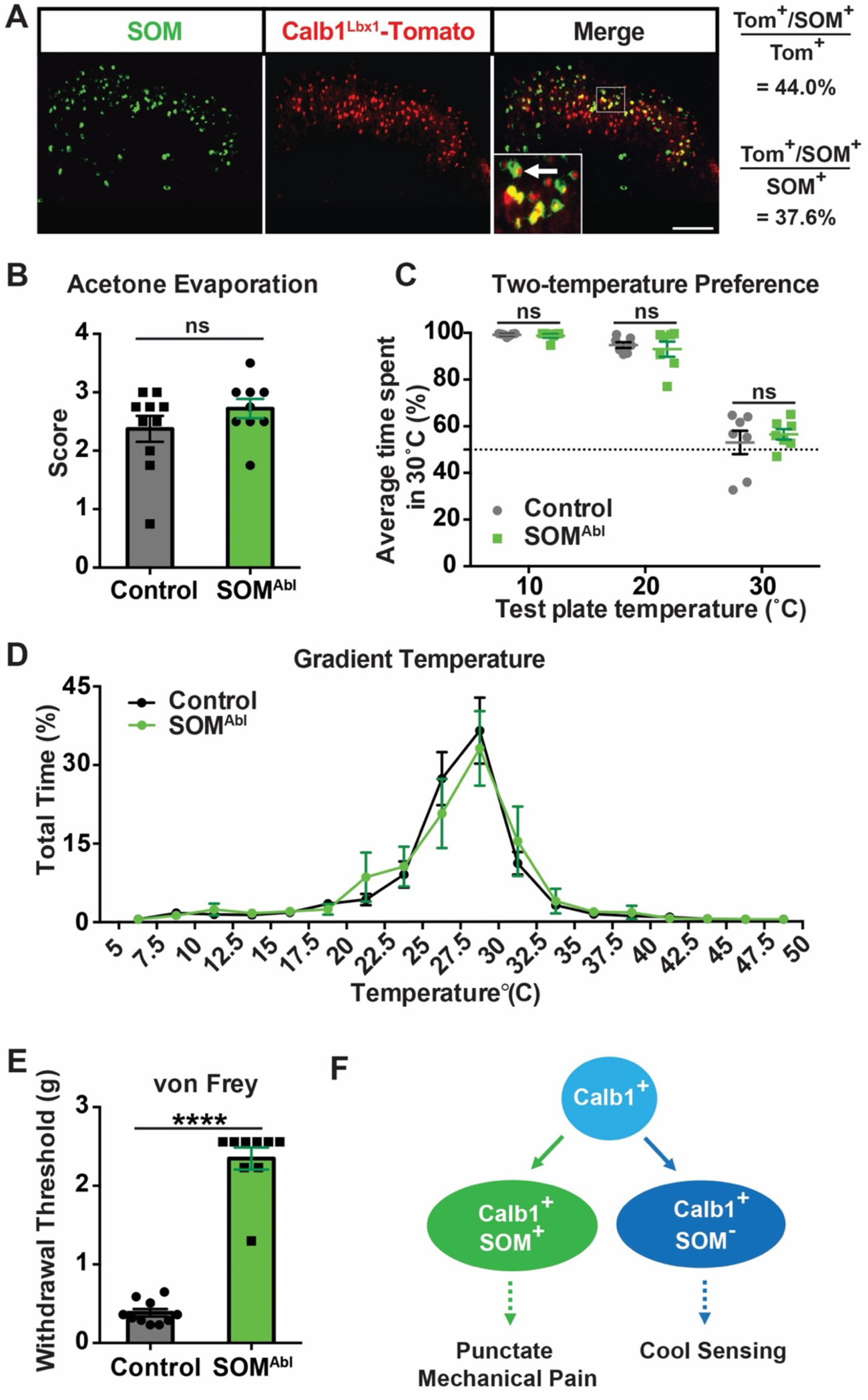
Ablation of SOM^Lbx1^ Neurons Leads to Deficits in Acute Punctate Mechanical Pain but Not Innocuous Cool Sensations. **(A)** Double staining of SOM mRNA (green) by *in situ* hybridization with Calb1^Lbx1^-Tomato signals (red). The percentage is calculated as double-positive neurons over total number of Calb1^Lbx1^-Tomato^+^ neurons (top) or double-positive neurons over total number of SOM^+^ neurons (bottom). Inset represents higher magnification of the boxed area. Arrow indicates double-positive cells for SOM and Tomato. n = 18 sections. Scale bar, 100 μm. **(B)** The acetone evaporation assay in SOM^Abl^ and control groups. Control: n = 10; SOM^Abl^: n = 9; ns, no significant difference; Mann-Whitney test. **(C)** Two-temperature preference between two temperature plates. Reference plate is set at 30°C, and test plate temperature is set at 30 °C, 20 °C and 10 °C. n = 7 in each group; ns, no significant difference; two-way ANOVA with Sidak post hoc analysis. Data points represent the average percentage of time spent on the reference plate across two trials over the total trial time. **(D)** Gradient temperature ranging from 5 °C to 50 °C was quantified as time spent in each temperature zone. Control: n = 7; SOM^Abl^: n = 5; ns, no significant differences; two-way ANOVA with Bonferroni post hoc analysis. **(E)** Acute punctate mechanical pain measured using the up-down von Frey assay. Control: n = 10; SOM^Abl^: n = 9; **** p < 0.0001; Student’s unpaired t test. **(F)** Schematic showing proposed Calb1^Lbx1^ subpopulations for cool sensations and acute punctate mechanical pain. Calb1^+^ neurons (light blue) represent the entire Calb1^Lbx1^ population, which can be further classified into at least two distinct subgroups: Calb1^Lbx1^;SOM^+^ (green) for mechanical punctate pain and Calb1^Lbx1^;SOM^-^ (dark blue) for cool sensing.

### Calb1^+^ neurons in the superficial dorsal horn receive inputs from TRPM8^+^ primary sensory neurons

Previous studies have shown that the TRPM8 channel, a non-selective cation channel, is sensitive to innocuous cool stimuli (McKemy et al., 2002; Peier et al., 2002). We asked whether TRPM8^+^ neurons in the DRG are the primary sensory neurons that provide inputs to Calb1^Lbx1^ neurons in the spinal cord. For this, we injected the TRPM8 agonist icilin into the hindpaw and recorded cool-induced (wet-dog shaking) behavioral responses. Compared to control littermates, Calb1^Abl^ mice showed nearly abolished wet-dog shaking responses (Figure 5A), but no change in acute nocifensive behaviors due to the intraplantar injection (Figure S6A). To further investigate the connection between TRPM8^+^ sensory neurons and Calb1^+^ neurons in the dorsal horn at a synaptic level, we crossed *Calb1^Cre^* and *TRPM8^GFP^* mice, then retro-orbitally injected *Calb1^Cre^*; *TRPM8^GFP^* mice with a *Cre*-dependent AAV-PHP.eB Brainbow virus that can stochastically express either teal fluorescent protein (TFP) or mCherry (mCh) fluorophores. This allowed us to sparsely label Calb1^+^ neurons in the spinal cord (hereafter referred to as Calb1^Brainbow^) without infecting the DRG (Figure S6B). Expansion microscopy was then utilized to obtain the nano-scale resolution necessary for synaptic identification without the use of super-resolution light microscopy (Figure 5B and 5C). Colocalization of pre- and post-synaptic markers, Bassoon and Homer, respectively, with Calb1^Brainbow^ and TRPM8^GFP^ were used to identify monosynaptic connections between TRPM8^+^ cooling inputs and Calb1^+^ spinal neurons (Figure 5C). Quadruple-positive synaptic interactions were identified predominantly in the TRPM8-innervation zone located in lamina I-II_o_ (Figure 5D and 5E), with unique multi-synaptic clusters identified (Figure S6C), suggesting that TRPM8^+^ cooling fibers synapse onto Calb1^+^ neurons in the superficial dorsal horn of the spinal cord. The monosynaptic connections from TRPM8^+^ sensory neurons to Calb1^+^ neurons were further confirmed by a pseudotyped rabies virus-based retrograde tracing combined with RNAscope staining in *Calb1^Cre^;Lbx1^Flpo^;Rosa26^ds-HTB^* mice (Figure S6D-S6F).

**Figure 5.**
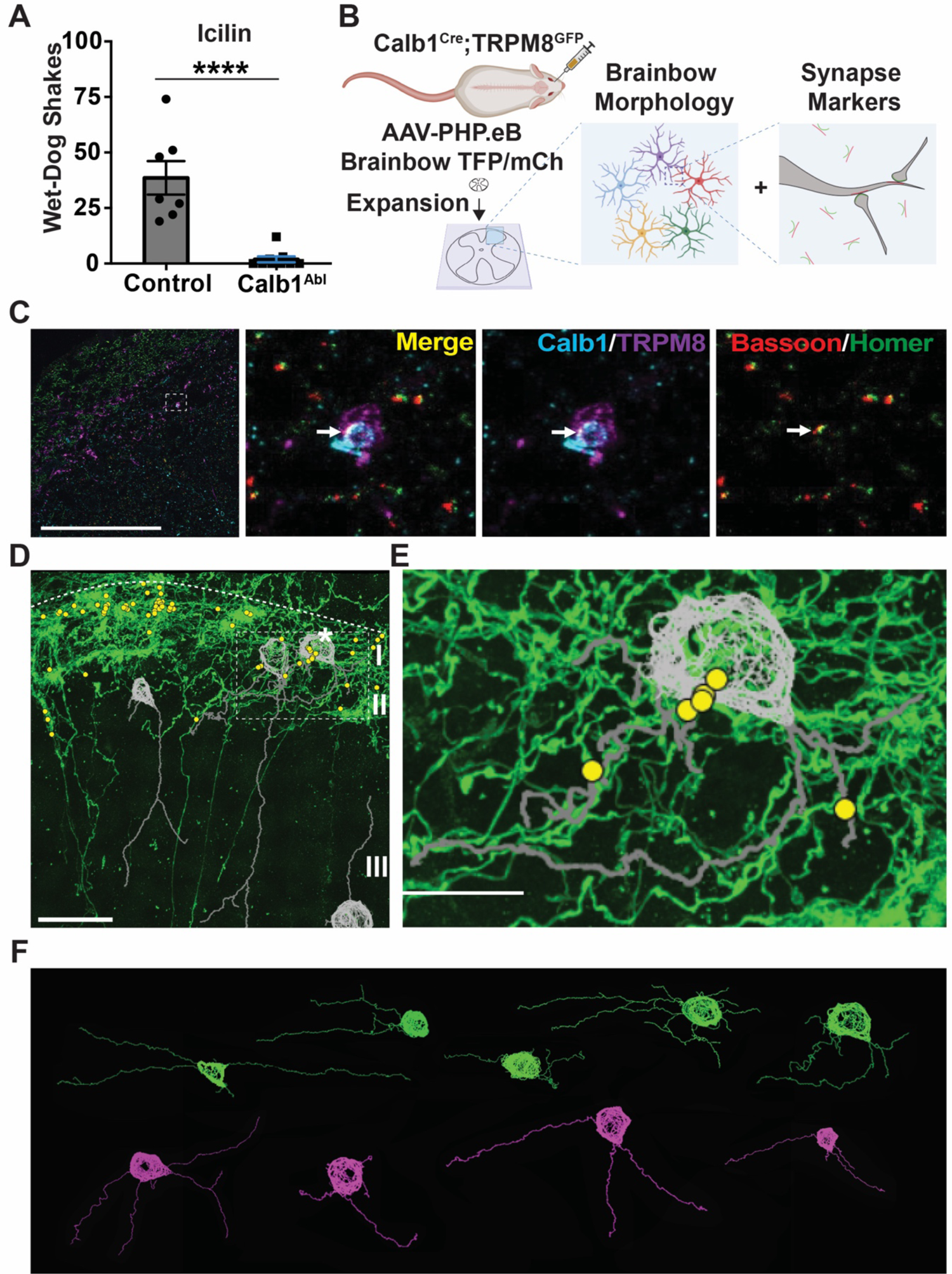
Calb1^+^ Neurons Receive Monosynaptic Inputs from Cool-sensitive TRPM8^+^ Primary Sensory Neurons in the Superficial Dorsal Horn of the Spinal Cord. **(A)** The number of wet-dog shakes in response to an agonist of TRPM8 channels, icilin, is abolished in Calb1^Abl^ mice compared to controls. Control: n = 7; Calb1^Abl^: n = 9; **** p < 0.0001; Student unpaired t test. **(B)** Schematic representing the experimental approach to virally label individual Calb1^+^ neurons in the spinal cord and TRPM8^GFP^ afferent neurons in the DRG using *Calb1^Cre^;TRPM8^GFP^* mice. Spinal cord tissue is embedded in a gel and expanded to enable reconstruction of morphology of Calb1^Brainbow^ neurons and identify the location of synaptic pairs. **(C)** Left: Overview of the dorsal horn of *Calb1^Cre^;TRPM8^GFP^* mice following Brainbow labeling and immunostaining excitatory pre- and post-synaptic markers Bassoon and Homer, respectively. Right three panels: Higher magnification of insets depicting representative images of quadruple-positive interaction. Cyan: The dendritic branch of a Calb1^Brainbow^ neuron; purple: the axon terminal of a TRPM8^+^ sensory neuron; red: the presynaptic marker Bossoon; green: the postsynaptic marker Homer. Arrow shows a quadruple-positive synaptic connection. Scale bar, 30 μm estimated based on anticipated expansion factor. **(D)** Representative image showing the location of identified quadruple-positive synaptic connections (yellow dots) and cell morphology of Calb1^Brainbow^ neurons (grey) across the superficial dorsal horn. Scale bar, 30 μm estimated based on anticipated expansion factor. **(E)** Higher magnification of inset depicting representative image showing the morphology of a Calb1^Brainbow^ neuron that forms synaptic pairs with TRPM8^+^ afferents. Green: TRPM8^GFP^; grey: Calb1^Brainbow^ neuron; yellow: identified synaptic pairs between TRPM8^+^ afferents and this Calb1^+^ neurons. Scale bar, 15 μm estimated based on anticipated expansion factor. **(F)** Schematic summarizing the morphology of Calb1^Brainbow^ neurons that form synaptic connections with TRPM8^+^ primary sensory neurons. Green: Calb1^Brainbow^ neurons with local dendritic arborization in TRPM8-innervation zone; magenta: Calb1^Brainbow^ neurons that contain at least one dendritic arbor outside of the TRPM8-innervating zone. n = 6 sections.

Cooling-sensitive neurons in lamina I were previously described as pyramid cells, while polymodal heat-pinch-cold neurons were described as multipolar cells (Han et al., 1998). The sparse labeling of Calb1^+^ neurons in the dorsal horn allowed us to identify the morphology of Calb1^Brainbow^ neurons that received TRPM8^+^ inputs (Figure 5D-5F). We found that TRPM8-innervating Calb1^Brainbow^ neurons exhibit diverse morphologies, with somas located in the TRPM8-innervation zone (lamina I-II_o_) (Figure 5F). While the majority of dendrites were located in lamina I-II_o_ (green neurons in Figure 5F), several examples of neurons with at least one dendritic branch outside of lamina I-II_o_ were identified (magenta neurons in Figure 5F), suggesting those neurons may also receive inputs from deeper layers of the dorsal horn.

### Two distinct Calb1^Lbx1^ subpopulations in the superficial dorsal horn

To further characterize the Calb1^Lbx1^ spinal subpopulations and their sensory inputs, we performed whole-cell patch clamp recordings from Tomato^+^ spinal neurons across laminae I-II in naïve *Calb1^Lbx1^*;Ai65 mice (Figure 6A and 6C), and then performed RT-PCR analysis to determine the type of neurons from which recordings were performed (Calb1^Lbx1^;SOM^+^ or Calb1^Lbx1^;SOM^-^; Figure S8A and S8B). As a starting point, we characterized the firing pattern of recorded neurons and found that the firing pattern was similar between both populations, with the majority displaying an initial bursting pattern (Figure 6B and 6D). Next, we used high-frequency stimulation to characterize the monosynaptic/polysynaptic inputs to Calb1^Lbx1^ neurons. Evoked excitatory postsynaptic currents (eEPSCs) and action potentials (APs) were recorded under normal condition or under disinhibition conditions with the presence of strychnine and bicuculline (Str & Bic) to block glycine receptors and GABA_A_ receptors, respectively. We found that 80.0% (28/35) of Calb1^Lbx1^;SOM^-^ neurons in lamina I-II_o_ received C fiber inputs under normal condition, 68.6% (24/35) of Calb1^Lbx1^;SOM^-^ neurons received monosynaptic C fiber inputs, and 54.3% (19/35) of Calb1^Lbx1^;SOM^-^ neurons generated APs (Figure 6E and S7H). Under disinhibition condition, 75.0% (24/32) of Calb1^Lbx1^;SOM^-^ neurons received monosynaptic C fiber and 71.9% (23/32) of Calb1^Lbx1^;SOM^-^ neurons generated APs (Figure 6E and S7H). By contrast, none of Calb1^Lbx1^;SOM^-^ neurons in lamina I-II_o_ received A*β* fiber inputs and very few receive A*δ* fiber inputs, even under disinhibition condition (Figure 6E, S7F and S7G).

**Figure 6.**
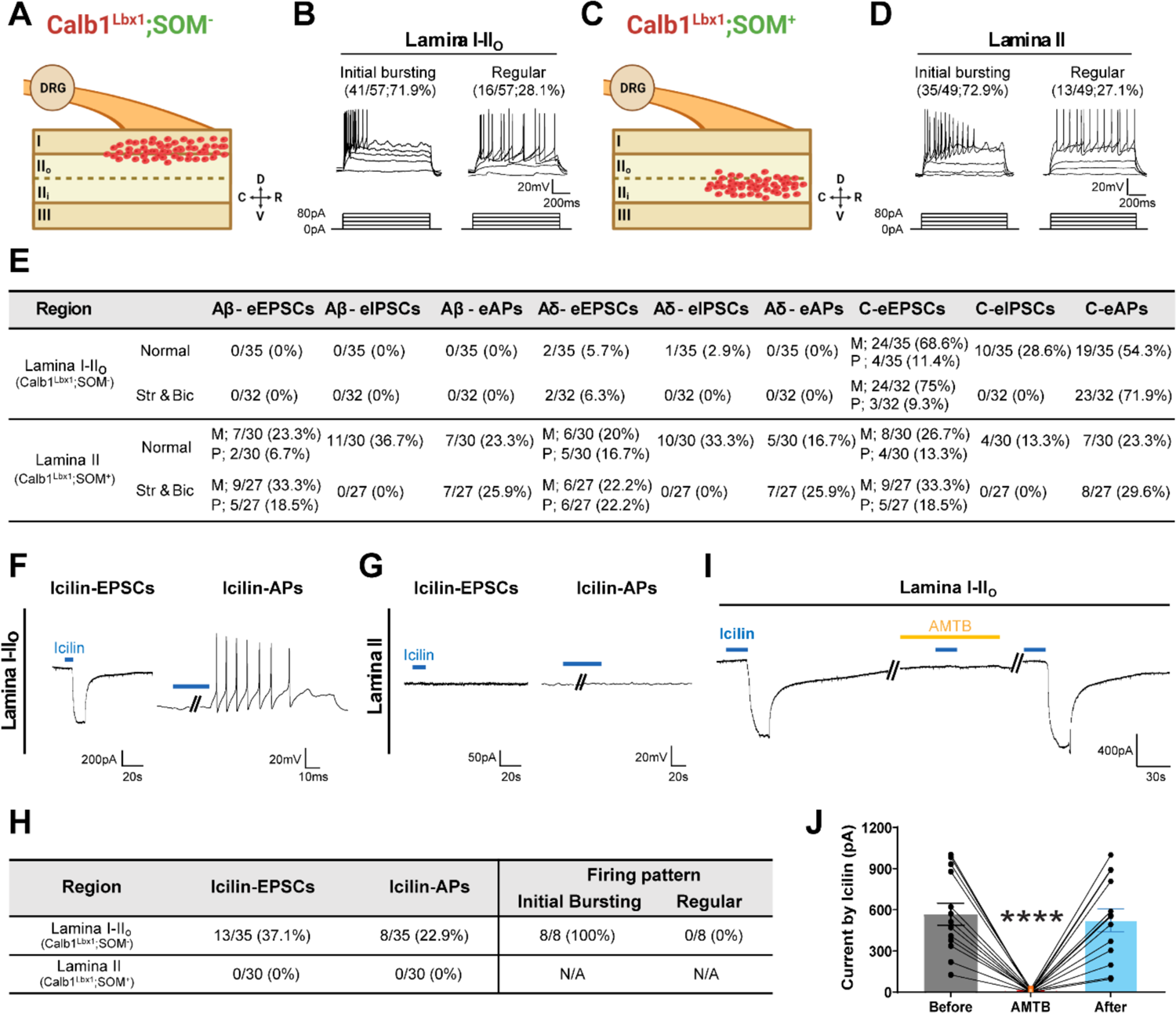
Firing Patterns and Sensory Inputs of Two Subpopulations of Calb1^Lbx1^ Neurons in the Superficial Dorsal Horn of the Spinal Cord. **(A)** Schematic demonstrating the location of Calb1^Lbx1^;SOM^-^ neurons in lamina I-II_o_. Red dots: recorded Calb1^Lbx1^;SOM^-^ neurons. n = 57 from 10 naïve *Calb1^Lbx1^*;Ai65 mice. **(B)** Firing properties of Calb1^Lbx1^;SOM^-^ neurons in lamina I-II_o_. **(C)** Schematic demonstrating the location of Calb1^Lbx1^;SOM^+^ neurons in lamina II. Red dots: recorded Calb1^Lbx1^;SOM^+^ neurons. n = 47 from 10 naïve *Calb1^Lbx1^*;Ai65 mice. **(D)** Firing properties of Calb1^Lbx1^;SOM^+^ neurons in lamina II. **(E)** Summary of A*β*-, A*δ*- or C-evoked EPSCs, IPSCs, and APs in Calb1^Lbx1^;SOM^-^ neurons in lamina I-II_o_ (top) and Calb1^Lbx1^;SOM^+^ neurons in lamina II (bottom) under normal conditions, and after strychnine and bicuculline application. Table represents a summary of sensory inputs in 35 Calb1^Lbx1^;SOM^-^ neurons in lamina I-II_o_ under normal condition, and 32 Calb1^Lbx1^;SOM^-^ neurons in lamina I-II_o_ upon strychnine and bicuculline application; 30 Calb1^Lbx1^;SOM^+^ neurons in lamina II under normal condition, and 27 Calb1^Lbx1^;SOM^+^ neurons in lamina II upon strychnine and bicuculline application. M, monosynaptic inputs. P, polysynaptic inputs. **(F)** Representative traces of icilin-induced eEPSC (left) and icilin-induced eAPs (right) in Calb1^Lbx1^;SOM^-^ neurons in lamina I-II_o_. Horizontal blue line indicates 1μM icilin application to the DRG chamber. **(G)** Representative traces of icilin-induced eEPSC (left) and icilin-induced eAPs (right) in Calb1^Lbx1^;SOM^+^ neurons in lamina II. Horizontal blue line indicates 1μM icilin application to the DRG chamber. **(H)** Summarized table of icilin-induced eEPSCs and icilin-induced eAPs at 1 μM concentration in Calb1^Lbx1^;SOM^-^ neurons in lamina I-II_o_ (top) and Calb1^Lbx1^;SOM^+^ neurons in lamina II (bottom) from 10 naïve *Calb1^Lbx1^*;Ai65 mice. Right panel: All icilin-responsive Calb1^Lbx1^;SOM^-^ neurons demonstrate an initial bursting firing pattern. **(I)** Representative trace showing a Calb1^Lbx1^;SOM^-^ neuron recorded upon administration of icilin (before), during co-administration of icilin and AMTB (middle), and upon administration of icilin (after). Diagonal lines indicate passage of time between chemical administrations. Blue lines represent 1 μM icilin application. Yellow line represents 100 μM AMTB administration. **(J)** Quantification of icilin-induced eEPSCs in Calb1^Lbx1^;SOM^-^ neurons. n = 13 from 10 naïve *Calb1^Lbx1^*;Ai65 mice; **** p < 0.0001 two-way ANOVA with Tukey post hoc analysis.

Most TRPM8^+^ primary sensory neurons are unmyelinated C nociceptors and could mediate the C-fiber evoked APs. Application of icilin to acutely disassociated TRPM8^GFP^ DRG neurons evoked Ca^2+^ influx in a dose dependent manner, with all neurons responding beginning at 1 μM, thereby mimicking cool-temperature induced activation of TRPM8^GFP^ DRG neurons (Figure S7A-S7C). To examine whether TRPM8^+^ C fibers activate Calb1^Lbx1^;SOM^-^ neurons in lamina I-II_o_, a specialized two-chamber recording apparatus was used to isolate the intact and attached DRG in a separate chamber from the spinal cord (Figure S7D). Application of icilin (1 μM) to the DRG chamber but not the spinal cord chamber elicits large EPSCs (Icilin-EPSCs or I-EPSCs) and AP responses (Icilin-APs or I-APs) in neurons located in lamina I-II_o_ (Figure S7E). We then recorded from Calb1^Lbx1^;SOM^-^ neurons in lamina I-II_o_ using icilin application to the DRG chamber. We found that 37.1% (13/35) of Calb1^Lbx1^;SOM^-^ neurons in lamina I-II_o_ generated icilin-evoked EPSCs (Figure 6F and 6H), all of which were blocked by co-application of TRPM8 antagonist, AMTB, to the DRG chamber (Figure 6I and 6J), suggesting that the Calb1^Lbx1^;SOM^-^ subpopulation receives C fiber inputs from TRPM8^+^ primary sensory neurons. Interestingly, 8 out of 13 icilin-responding neurons generated APs (Figure 6H), and all exhibited initial bursting firing pattern (Figure 6H) and received monosynaptic C fiber-inputs (Figure S8A). These results indicate the presence of a unique subpopulation of Calb1^Lbx1^;SOM^-^ neurons in lamina I-II_o_ is cooling-sensitive.

Next, we examined the sensory inputs to Calb1^Lbx1^;SOM^+^ neurons in lamina I-II. We found that few recorded Calb1^Lbx1^ neurons in lamina I were SOM^+^ and the majority of Calb1^Lbx1^;SOM^+^ neurons were located in lamina II_i_ (Figure 6C). In total, 30 Calb1^Lbx1^;SOM^+^ neurons in lamina II were recorded under normal conditions, 40.0% (12/30) of Calb1^Lbx1^;SOM^+^ neurons received C fiber inputs and 23.3% (7/30) of Calb1^Lbx1^;SOM^+^ neurons generated AP outputs (Figure 6E, S7K and S8B). Under disinhibition condition, 51.8% (14/27) of Calb1^Lbx1^;SOM^+^ neurons received C fiber inputs and 29.6% (8/27) of Calb1^Lbx1^;SOM^+^ neurons generated AP outputs (Figure 6E, S7K and S8B). In addition, we found that about half of Calb1^Lbx1^;SOM^+^ neurons in lamina II received A*β* fiber and (or) A*δ* inputs (Figure 6E, S7I, S7J and S8B). Interestingly, we found that none of Calb1^Lbx1^;SOM^+^ neurons in lamina II generated icilin-EPSCs or icilin-APs (Figure 6G, 6H and S8B), suggesting that Calb1^Lbx1^;SOM^+^ neurons do not receive inputs from TRPM8^+^ primary sensory neurons. Given the loss of acute punctate mechanical pain in SOM^Abl^ and Calb1^Abl^ mice, C fiber sensory neurons that form synaptic connections with Calb1^Lbx1^;SOM^+^ neurons in lamina II likely represent mechanical nociceptors.

### Calb1^Lbx1^ interneurons amplify the activity of cooling-sensitive SPB neurons

In lamina I, ∼95% of spinal projection neurons target the lateral parabrachial nucleus (Todd, 2010). To determine whether Calb1^Lbx1^ spinal neurons that transmit cool sensations are interneurons or projection neurons, we first bilaterally injected fluorophore-conjugated cholera toxin subunit-B (CTB) into the lateral parabrachial nucleus followed by acetone administration onto the hindpaw to induce c-Fos expression in the ipsilateral spinal cord (Figure 7A and 7B). Although double-positive neurons (CTB^+^ and Fos^+^) representing cool-sensitive SPB neurons were detected, none of the cooling-sensitive SPB neurons were Tomato^+^ in *Calb1^Lbx1^*;Ai65 mice (Figure 7C), confirming that Calb1^Lbx1^ neurons are local interneurons and not cooling-sensitive SPB neurons.

**Figure 7.**
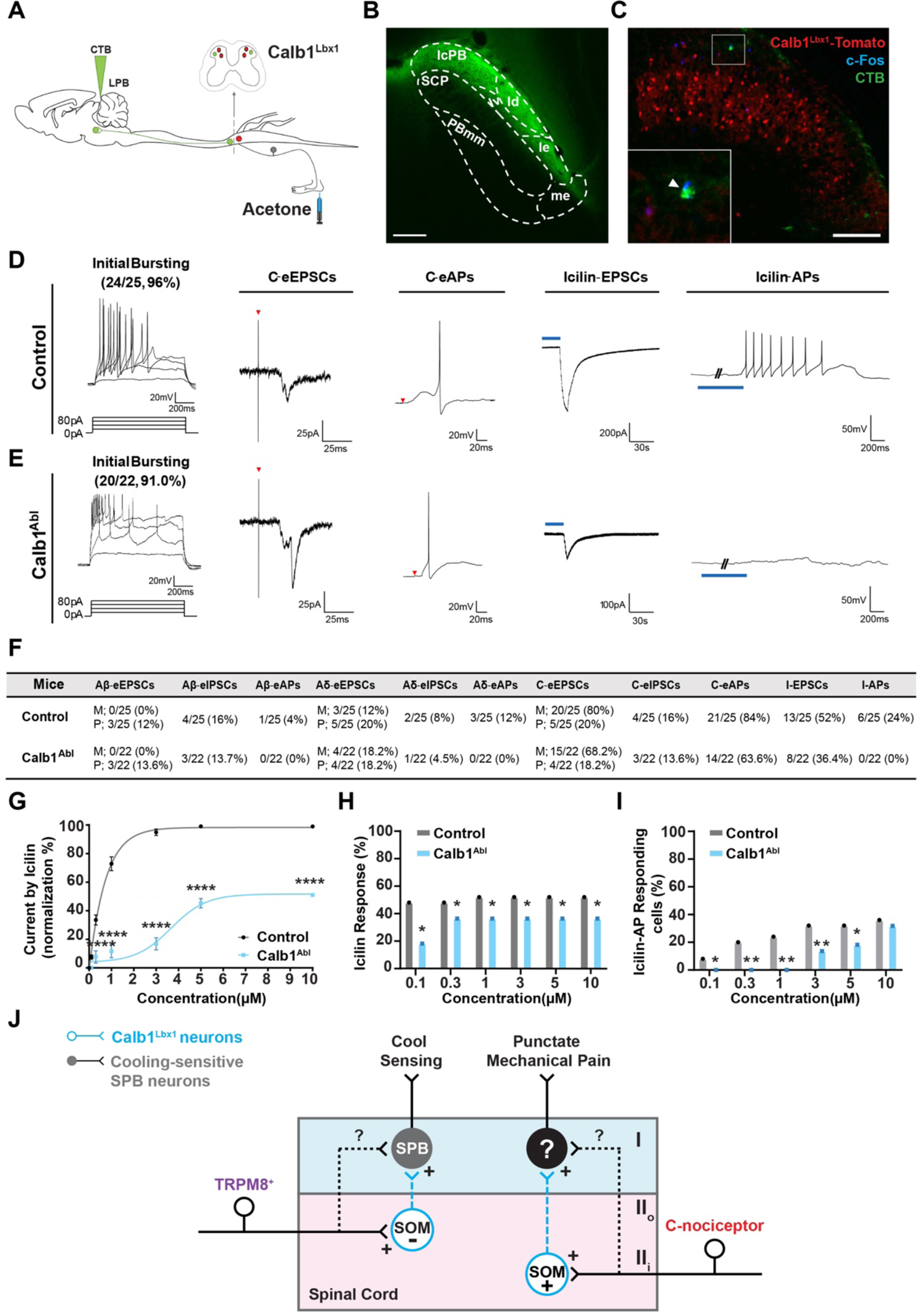
Calb1^Lbx1^ Neurons Amplify the Activity of Cooling-sensitive SPB Neurons. **(A)** Schematic demonstrating bilateral injection of fluorescent fluorophore conjugated cholera toxin subunit B (CTB) into the lateral parabrachial nucleus to label SPB neurons (green) and acetone co-administration onto hindpaw to label cooling-sensitive neurons in *Calb1^Lbx1^*;Ai65 mice (red). **(B)** Representative image of the injection site in the lateral parabrachial nucleus. Scale bar, 100 μm. **(C)** Out of 68 labeled SPB neurons, no triple-positive (red: Calb1^Lbx1^-Tomato^+^; green: CTB; blue: Fos^+^) cool SPB neurons were identified (0%, 0/68) in the ipsilateral dorsal horn of the spinal cord following acetone treatment. Inset represents high magnification of the boxed area. Arrowhead indicates a c-Fos and CTB double-positive cooling-sensitive SPB neuron that is Tomato^-^. n = 9 sections from 3 mice. Scale bar, 100 μm. **(D-E)** Representative firing properties, C fiber inputs, and icilin-induced eEPSCs and eAPs in CTB^+^ SPB neurons in control **(D)** and Calb1^Abl^ **(E)** mice. The majority of CTB^+^ SPB neurons in control (96%, 24/25) and Calb1^Abl^ (91.0%, 20/22) animals display an initial bursting firing pattern upon current injection. Horizontal blue line: Icilin 1μM. Red triangle: presentation of stimulus. **(F)** Summary of A*β*-, A*δ*-, and C-evoked EPSC, IPSC, and APs, and icilin-induced eEPSCs and APs in CTB^+^ SPB neurons in control and Calb1^Abl^ mice. Control: 25 neurons from 5 mice; Calb1^Abl^: 22 neurons from 4 mice. M, monosynaptic inputs. P, polysynaptic inputs. Icilin: 1 μM. **(G)** Dose response curve of icilin (μM)-induced eEPSCs in CTB^+^ SPB neurons in control and Calb1^Abl^ mice. Control: 13 neurons from 5 mice; Calb1^Abl^: 8 neurons from 4 mice; **** p < 0.0001; *** p < 0.001; ** p < 0.01; * p < 0.05; two-way ANOVA with Sidak post hoc analysis. **(H)** Quantification of icilin-evoked EPSCs in CTB^+^ SPB neurons at different concentrations in control and Calb1^Abl^ mice. Control: 13 neurons from 5 mice; Calb1^Abl^: 8 neurons from 4 mice; * p < 0.05; two-way ANOVA with Tukey post hoc analysis. **(I)** Quantification of icilin-evoked APs in CTB^+^ SPB neurons at different concentrations in Control and Calb1^Abl^ mice. Control: 13 neurons from 5 mice; Calb1^Abl^: 8 neurons from 4 mice; ** p < 0.01; * p < 0.05; two-way ANOVA with Tukey post hoc analysis. **(J)** Schematic showing proposed neural pathways that transmits innocuous cool sensations and acute punctate mechanical pain. Calb1^Lbx1^;SOM^-^ interneurons in laminae I-II_o_ receive monosynaptic inputs from TRPM8^+^ primary sensory neurons and then innervate to cooling-sensitive SPB neurons via monosynaptic or polysynaptic connections; whereas Calb1^+^;SOM^+^ interneurons in lamina II are proposed to receive inputs from mechanosensitive C-nociceptors and then connect to unknown SPB neurons for acute punctate mechanical pain. However, whether TRPM8^+^ fibers or C-nociceptor fibers synapse onto SPB neurons in lamina I remains unknown.

To examine the function of Calb1^Lbx1^ neurons in the cooling-transmission pathway, we recorded from CTB-labeled SPB neurons in control and Calb1^Abl^ naïve mice using the two-chamber recording system (Figure 7D). Icilin application to the DRG chamber was used to identify cooling-sensitive SPB neurons. We found that the vast majority of SPB neurons exhibit an initial bursting pattern in both groups (Figure 7D and 7E). Next, we examined the sensory inputs to recorded SPB neurons. In control mice, all recorded SPB neurons in lamina II_o_ received C fiber inputs and 84.0% (21/25) of SPB neurons generated APs (Figure 7D and 7F). In Calb1^Abl^ mice, 86.4% (19/22) of recorded SPB neurons received C fiber inputs and 63.6% (14/22) of recorded SPB neurons generated APs (Figure 7E and 7F). Compared to control animals, ∼26% (1-63.6/86.4) of SPB neurons were not activated by C fiber inputs after ablating Calb1^Lbx1^ neurons. We also detected a small portion of SPB neurons that received polysynaptic A*β*inputs and mono/polysynaptic A*δ* inputs in control mice (Figure 7F). However, A*β*/A*δ*evoked APs were abolished in Calb1^Abl^ mice, suggesting a small population of Calb1^Lbx1^ neurons may link A*β*/A*δ* inputs with SPB neurons. Next, we recorded icilin (1 μM)-evoked EPSCs and APs in cooling-sensitive SPB neurons. We found that icilin-evoked EPSCs and APs in 52.0% (13/25) and 24.0% (6/25) of SPB neurons in control mice, respectively (Figure 7D and 7F). By contrast, a small portion of icilin-evoked EPSCs (36.4%, 8/22) and no icilin-evoked APs (0%, 0/22) were recorded in SPB neurons of Calb1^Abl^ mice (Figure 7E and 7F), consistent with the reduction of C fiber-evoked EPSCs and APs in Calb1^Abl^ mice. Interestingly, cooling-sensitive SPB neurons display a right-shifted icilin-EPSCs dose response curve in Calb1^Abl^ mice compared to controls (Figure 7G) with significance reductions in icilin-EPSCs (Figure 7H) and icilin-APs (Figure 7I) at different concentrations of icilin. These results suggest that Calb1^Lbx1^ spinal interneurons are essential for the activation of cooling-sensitive SPB neurons, opening up the possibility that Calb1^Lbx1^ interneurons may act as a cool-signal amplifier to prioritize cool temperature information for projection neuron integration in the spinal cord (Figure 7J).

## DISCUSSION

In the present study, we demonstrated the presence of a subpopulation of Calb1^Lbx1^ excitatory interneurons in the superficial spinal cord that is essential for innocuous cool temperature transmission, but not noxious cold. Particularly, we identified at least two functionally distinct subpopulations of Calb1^Lbx1^ neurons in lamina I-II of the dorsal horn of the spinal cord: 1) a Calb1^Lbx1^;SOM^-^ subpopulation of excitatory interneurons in lamina I-II_o_ that transmits cool sensations, and 2) a Calb1^Lbx1^;SOM^+^ subpopulation of excitatory interneurons in lamina II that transmits acute punctate mechanical pain. In lamina I-II_o_, most Calb1^Lbx1^ neurons are SOM^-^ and receive C-fiber inputs. Cooling-activated Calb1^+^ neurons cover ∼50% Calb1^Lbx1^;SOM^-^ neurons in lamina I-II_o_ (Figure 6E). Lamina I-II_o_ Calb1^Lbx1^;SOM^-^ neurons are innervated by TRPM8^+^ C-cooling neurons and amplify cooling signals to projection neurons in lamina I. However, we do not exclude the possibility that the TRPM8-innervated Calb1^+^ neurons may be polymodal.

In lamina II, most neurons receive C and A*δ* inputs directly and some neurons also receive polysynaptic inputs from A*β* fibers (Braz et al., 2014; Todd, 2010). MrgprD^+^ polymodal nociceptors innervate multiple types of interneurons in lamina II (Wang and Zylka, 2009). Interestingly, ablation of MrgprD^+^ nociceptors or SOM^+^ interneurons in the dorsal horn of the spinal cord attenuated acute punctate mechanical pain (Cavanaugh et al., 2009; Duan et al., 2014). Thus MrgprD^+^ nociceptors may connect with Calb1^+^;SOM^+^ interneurons for transmitting acute punctate mechanical pain. Previous single-cell RNA sequence results showed that Calb1 partially overlaps with CCK and neurotensin (NTs) in lamina III (Haring et al., 2018; Russ et al., 2021). Ablation of CCK^+^ neurons impaired brush-induced paw withdrawal responses and activation of NTs^+^ neurons in lamina III facilitated brush-induced paw withdrawal responses (Gatto et al., 2021). Our results suggest that Calb1^+^;CCK^+^;NTs^+^ interneurons in lamina III may be required for transmitting innocuous paw withdrawal responses.

Previous *in vivo* single unit recording studies showed that an innocuous cool stimulus can activate two groups of spinal neurons: (1) “COOL” neurons responding selectively to innocuous cool, and (2) CMH neurons responding to innocuous cool, noxious mechanical (pinch) and heat stimuli (Craig et al., 2001). Consistently, *in vivo* imaging results in lamina I SPB neurons confirmed that there are two populations of cooling-sensitive neurons: cooling-specific projection neurons (14%) and polymodal projection neurons that response to cool, cold, pinch and heat (76%) (Chisholm et al., 2021). Under this paradigm it is possible that distinct spinal populations for cold/cool sensing exist, in particular, a population specific for innocuous cool sensation and a population of polymodal (combination of cold/cool, mechanical pain, and/or heat pain) neurons for burning sensation. Lamina I projection neurons receive monosynaptic connections from the DRG (Grudt and Perl, 2002) and interneurons in lamina I and II (Luz et al., 2010). Our present study reveals a feed-forward microcircuit that transmits innocuous cool sensations in the superficial dorsal horn. In this circuit, a small cluster of Calb1^Lbx1^;SOM^-^ neurons (∼23%, 8/35, Figure 6H) in lamina I-II_o_, exhibits an initiate bursting firing pattern and receives monosynaptic inputs from TRPM8^+^ cooling-sensitive primary sensory neurons.

Activation of TRPM8^+^ neurons is able to evoke AP firing in Calb1^Lbx1^;SOM^-^ neurons, and pharmacological silencing of Calb1^Lbx1^ neurons largely attenuated the activity of cooling-sensitive SPB neurons. There results suggest that the Calb1^Lbx1^;SOM^-^ subpopulation of neurons presents a potent source of excitatory signaling that could amplify cooling afferent outputs to cooling-sensitive SPB neurons. Further studies characterizing the synaptic connections from Calb1^Lbx1^;SOM^-^ neurons to cooling-sensitive SPB neurons will advance our understanding of the role in cool transmission.

Taken together, our present study has identified a small cluster of Calb1^Lbx1^ excitatory interneurons in the superficial dorsal spinal cord that acts as a critical node in a circuit linking cooling-sensitive TRPM8^+^ primary sensory neurons and SPB neurons for innocuous cooling transmission from the skin to the brain.

## ACKNOWLEDGMENTS

We thank Dr. Martyn Goulding for *Lbx1^Flpo^*, *Tau^ds-DTR^*, and *Rosa26^ds-HTB^* mice, Dr. Susan Dymecki for *Rosa26^CAG-ds-hM4Di^* mice, Dr. Hongkui Zeng for Ai14 and Ai65 mice, and Dr. Karel Svoboda for *Rosa26^CAG-ds-ReaChR^*. We thank Drs. Zhigang He, Chen Wang, and Yuanyuan Liu for sharing EnvA-pseudotyped, G-deleted-mCherry rabies virus. We are grateful to Dr. Mohammed Akaaboune for comments on an earlier version of the manuscript. We appreciate the encouragement and helpful comments from other members of the Duan laboratory. This work was supported by NINDS (5R01NS109170 to Duan; 1R01NS118769 to Xu and Duan), NIGMS (5R35GM126917 to Xu) and NIMH (1RF1MH120005 to Cai).

## AUTHOR CONTRIBUTIONS

B.D conceptualized and directed this project. L.H., H.L., C.C.H., I.R., and T.L.R.F. performed behavior experiments. H.L. performed electrophysiological recordings. L.H. performed histology studies. L.H., F.Y.S., E.P., and D.C. performed Brainbow staining, expansion microscopy, and data interpretation. W.C., H.L., and X.Z.S.Xu performed Ca^2+^ imaging and data interpretation. C.L. and K.P.P. constructed the gradient temperature apparatus. B.D. wrote the manuscript with L.H., H.L., and input of other authors.

## STAR★Methods

### KEY RESOURCES TABLE

| Reagent or Compound | Source | Identifier |
| --- | --- | --- |
| **Antibodies** |  |  |
| Rabbit anti-NK1R | MilliporeSigma | Cat#S8305; RRID: AB_261562 |
| Rabbit anti-α-CGRP | Peninsula Lab | Cat# T-4032; RRID: AB_2307330 |
| Alexa Fluor 647-conjugated Isolectin GS-IB4 | ThermoFisher | Cat#I32450; RRID: SCR_014365 |
| Rabbit anti-PKCγ | Santa Cruz | Cat# sc-211; RRID: AB_632234 |
| Rabbit anti-Pax2 | ThermoFisher | Cat# 71-6000; RRID: AB_2336046 |
| Rabbit anti c-Fos | Abcam | Cat# ab190289, RRID:AB_2737414 |
| Goat anti-Rabbit IgG (H+L) Alexa Fluor plus 647 | ThermoFisher | Cat# A32733; RRID: AB_2633282 |
| Goat anti-Mouse IgG (H+L) Alexa Fluor plus 488 | ThermoFisher | Cat# A32723; RRID: AB_2633275 |
| Goat anti-Rabbit IgG (H+L) Alexa Fluor plus 488 | ThermoFisher | Cat# A32731; RRID: AB_2633280 |
| Mouse anti-Calb1 | EnCor Biotech | Cat# MCA-5A9 RRID:AB_2572239 |
| Chicken anti-Homer | Synaptic Systems | Cat# 160 006 RRID:AB_2631222 |
| Guinea Pig anti-Bassoon | Synaptic Systems | Cat# 141 004 RRID:AB_2290619 |
| Rabbit anti-mCherry | Cai Lab custom made | NA |
| Rabbit anti-TFP | Cai Lab custom made | NA |
| Sheep anti-GFP | Biorad | Cat# 4745-1051 |
| Donkey anti-Sheep IgG (H+L) Alexa Fluor plus 488 | Jackson Immunoresearch | Cat# 713-546-147 RRID: AB_2340746 |
| Donkey anti-rabbit IgG (H+L) Alexa Fluor plus Cy3 | Jackson Immunoresearch | Cat# 711-166-152 RRID: AB_2313568 |
| Donkey anti guinea pig IgG (H+L) Alexa Fluor plus 594 | Jackson Immunoresearch | Cat# 706-585-148 RRID: AB_2340474 |
| Donkey anti chicken IgG (H+L) Alexa Fluor plus 647 | Jackson Immunoresearch | Cat# 703-606-155 RRID: AB_2340380 |
| **Bacterial and Viral Strains** |  |  |
| EnvA-pseudotyped G-deleted-mCherry rabies virus | Generated by Cai lab | Addgene #32636 |
| pAAVPHPeB-EF-mChT(Bb1.0) | Generated by Cai lab | NA |
| Cholera Toxin Subunit B (Recombinant), Alexa Fluor 488 Conjugate | ThermoFisher | Cat# C34775 |
| Cholera Toxin Subunit B (Recombinant), Alexa Fluor 647 Conjugate | ThermoFisher | Cat# C34778 |
| **Chemicals, Peptides and Recombinant Proteins** |  |  |
| Diphtheria toxin | Millipore Sigma | Cat# D0564 |
| Clozapine N-oxide (CNO) | Millipore Sigma | Cat# C0832 |
| Icilin | Millipore Sigma | Cat# I9532 |
| AMTB hydrochloride | Tocris | Cat# 3989 |
| Acetone | Millipore Sigma | Cat# 650501 |
| Strychnine | Millipore Sigma | Cat# S0532 |
| Bicuculline | Millipore Sigma | Cat# B7561 |
| **Experimental Models:**<br>**Organisms / Strains** |  |  |
| Mouse: B6;129S-<br>*Calb1<sup>tm2.1(cre)Hze</sup>/J* | Jackson Laboratory | Cat# 028532 |
| Mouse: *SOM-IRES-Cre* | Jackson Laboratory | Cat # 013044 |
| Mouse: *B6;129S-Lbx1 <tm1(flpo)Gou > (Lbx1-Flpo)* | Generated by Goulding Lab | Duan et al., 2014 |
| Mouse: *Rosa26<sup>CAG-LSL-FSF-HTB</sup>* | Generated by Goulding Lab | Bourane et al., 2015 |
| Mouse: B6.129-Tau<br><tm1(LSL.FSF.DTR)Gou>(Ta<br>u-loxP-STOP-loxP-FRT-STOP-FRT-DTR) | Generated by Goulding Lab | Duan et al., 2014 |
| Mouse: *B6;129S6-*<br>*Gt(ROSA)26Sortm9(CAG-*<br>*mCherry,- CHRM4\*)Dym/J*<br>*(Rosa26<sup>CAG-FSF-LSL-</sup>*<br>*<sup>hM4Di</sup>)* | Jackson Laboratory | Cat# 029040 |
| Mouse: *B6;129S-*<br>*Gt(ROSA)26Sortm2.1Ksvo/J* | Jackson Laboratory | Cat# 024846 |
| *TRPM8<sup>GFP</sup>* | Generated by Patapoutian Lab | Dhaka et al., 2007 |
| Mouse: Ai14 | Jackson Laboratory | 007914 |
| Mouse: Ai65 | Jackson Laboratory | 021875 |
| **Oligonucleotides** |  |  |
| GAPDH see table 1 | Primer-BLAST | N/A |
| SOM (Sst) see table 1 | Primer-BLAST | N/A |
| **Algorithms and Software** |  |  |
| Adobe Creative Cloud<br>(Illustrator / Photoshop /<br>Premiere Pro) | Adobe | https://www.adobe.com/ |
| ImageJ | NIH | https://imagej.nih.gov/ij/ |
| Prism 6, 9 | GraphPad Software | RRID: SCR_002798 |
| MATLAB R2020, 2021 | MathWorks | https://www.mathworks.com/products/matlab.html |
| MetaFluor | Molecular Devices | RRID: SCR_014294 |
| pClamp 10.0 | Molecular Devices | RRID: SCR_011323 |

### LEAD CONTACT AND MATERIALS AVAILABILITY

This study did not generate new unique reagents. Further information and requests for resources and reagents should be directed to and will be fulfilled by the Lead Contact, Dr. Bo Duan.

### EXPERIMENTAL MODEL AND SUBJECT DETAILS

All animal experiments were performed in accordance with protocols approved by the Institutional Animal Care and Use Committee at University of Michigan following NIH guidelines. Both male and female mice were used for all experiments. Mice were group housed at room temperature with *ad libitum* access to standard lab mouse pellet food and water on a 12 hours light/12 hours dark cycle. The mouse lines used in the present study were: *Calb1^Cre^* (#028532, JAX), *SOM^Cre^* (#013044, JAX), *Rosa26^LSL-tdTomato^* (Ai14, #007914, JAX), Ai65 (#021875, JAX), *Lbx1^Flpo^* (Duan et al., 2014), *Rosa26 ^CAG-ds-hM4Di^* (#029040, JAX), *Rosa26 ^CAG-ds-ReaChR^* (#024846, JAX), *Rosa26 ^CAG-LSL-FSF-HTB^* (*Rosa26^ds-HTB^*) (Bourane et al., 2015) and *Tau^LSL-FSF-DTR^* (*Tau^ds-DTR^*) (Duan et al., 2014). *SOM^Cre^* mice were crossed with Ai14 reporter mice (*SOM^Cre^*;Ai14) to label the SOM^Cre^-derived neurons. *Calb1^Cre^* mice were crossed with *Lbx1^Flpo^* and *Rosa26^CAG-ds-tdTomato^* (Ai65) reporter mice to label the *Calb1^cre^* and *Lbx1^Flpo^*-derived (Calb1^Lbx1^) neurons. We ablated DTR-expressing neurons as previously described (Duan et al., 2014). 6-10 weeks old mice were intraperitoneally injected with diphtheria toxin (DTX, 50 mg/kg; MilliporeSigma, St. Louis, MO) at day 1, day 4, and day 7. In most strains, we performed behavioral or histochemical experiments 4 weeks after DT injection. *Rosa26 ^CAG-ds-hM4Di^* mice were crossed with *Calb1^Cre^* mice and *Lbx1^Flpo^* mice, clozapine N-oxide (CNO, 5 mg/kg, MilliporeSigma, St. Louis, MO) was injected to acutely silence Calb1^+^ neurons. Behavioral tests were performed 40 minutes after CNO injection. Rectal temperature was assessed after all behavioral experiments were completed. *Calb1^Cre^* mice were crossed with *TRPM8^GFP^* mice and injected with AAV-PHP.eB-mCherry-TFP virus at 6-10 weeks old; 3 to 4 weeks later the mice were perfused for histology experiments.

## METHOD DETAILS

### *In Situ* Hybridization and Immunohistochemistry

*In situ* hybridization (ISH) procedures were performed to detect mRNA expression as described previously (Pan et al., 2019). Prior to performing ISH, Tomato fluorescent signals were first captured under a fluorescent microscope (Leica DMi8, Germany) for double staining analysis. After ISH, bright field images were converted into pseudo-fluorescent signals and merged onto the Tomato before images in Photoshop (Adobe Photoshop CS6). 3-5 mice per genotype were used for quantitative analysis. Only cells containing nuclei and showing levels of expression or staining clearly above background were scored. To detect protein expression, immunohistochemistry was performed using rabbit anti-NK1R (1:1000, #S8305 MilliporeSigma, St. Louis, MO), rabbit anti-*α*-CGRP (1:500, #T-4032, Peninsula Lab, San Carlos, CA), Alexa fluor 647-conjugated isolectin GS (IB4) (10 μg/mL, #I32450, ThermoFisher Scientific, Waltham, MA), rabbit anti-PKC*γ*(1:500, # sc-211, Santa Cruz Biotechnology, Dallas, TX), rabbit anti-Pax2 (1:100, #71-6000,ThermoFisher, Waltham, MA), mouse anti-Calb1 (1:500,# MCA-5A9, EnCor Biotech Gainesville, FL), or rabbit anti-c-Fos (1:500, #ab190289, Abcam, Cambridge, United Kingdom) which were diluted in 0.2% of Triton X-100 and 10% normal goat serum in PBS (blocking buffer) and photographed under a fluorescent microscope.

### RNAscope

To detect *TRPM8* mRNA colocalization with rabies virus-induced mCherry expression in presynaptic neurons, RNAscope was utilized. As described previously (Gong et al., 2019), paraformaldehyde(PFA)-fixed mouse DRG segments from L1 to L6 were frozen in optimal cutting temperature (OCT) freezing medium then cryosectioned (12-16 μm thickness) onto glass slides and stored at −20 °C. Prior to performing RNAscope, a fluorescent microscope (Leica DMi8, Germany) captured Tomato fluorescent signals. RNAscope was performed using a TRPM8-probe (1:2000, #420451-C3, ACD Bio, Newark, CA) with TSA 647 fluorophore (#K1052-100-300, Apex Bio, Houston, TX) according to manufacturer’s instructions. After RNAscope, Photoshop (Adobe Photoshop CS6) was used to merge fluorescent signals onto the Tomato before images.

### Behavioral Tests

Mice of either sex were used, and for all behavior tests the experimenter was blinded to the genotype of the animals and littermate mice (B6J/129 mixed genetic background) were used as controls. After three to five ‘habituation’ sessions (20 minutes per day) in the behavior testing apparatus, acute somatosensory measures were recorded on five consecutive days in the given order: rotarod, light brushing, von Frey and Hargreaves (day 1); hot plate and cold plate (day 2), acetone and sticky tape (day 3); pinprick and pinch (day 4), dry ice (day 5). A cutoff of 60 seconds (46 °C), 30 seconds (50 °C), 20 seconds (54 °C), and 15 seconds (pinch) was applied to prevent injury to the animal (Duan et al., 2014; Pan et al., 2019). Temperature gradient assay was performed over at least two days to allow for complete habituation before the testing began. Similarly, the two-temperature preference assay was performed over three to five days to prevent the development of a place preference (not a temperature preference).

### Two-Temperature Preference Assay

To test preference when given the choice between two temperatures (two-temperature preference), mice were placed onto two adjacent temperature plates (BIO-CHP Cold Hot Plate Test, Bioseb, Pinellas Park, FL) for five minutes and the time spent on each plate was recorded (BIO-T2CT, Bioseb, Pinellas Park, FL). If mice exhibited a place preference (less than 30% time spent on each plate), when both temperature plates were set to the same temperature (30 °C), they were excluded from further testing. To avoid the development of a place preference due to a negative association of the chamber to a noxious temperature, mice were tested over three days in the following order: 30 °C vs 30 °C and 30 °C vs 20 °C (day 1), 30 °C vs 10 °C (day 2), 30 °C vs 50 °C (day 3).

### Gradient Temperature Assay

The gradient temperature device is a rectangular plate machined out of copper 101, with the top surface deburred to allow for a uniform and smooth finish. Thermoelectric coolers (TECs) with heatsinks are installed at several places underneath the assay. The TECs are feedback-controlled and can operate in either cooling or heating mode, allowing for customized temperature profiles along the length of the plate. Mice were acclimatized in the arena for at least 30 minutes or until the mouse was habituated, followed by a 30-minute video recording period to track the mouse’s movement within the arena. Only one mouse was tested at a time. The gradient temperature arena measures 140 cm in length, 10 cm in width and has opaque plexiglass walls with a height of 40 cm. The copper floor of the arena was maintained at a gradient temperature of 5-50 °C. The arena is virtually divided into 18 zones with distinct temperatures. Time spent within each zone was analysed using MATLAB.

### Acetone Evaporation Assay

Animals were placed in an elevated chamber with mesh floor. During ‘habituating’ periods, acetone was exposed to the environment to control for olfactory stimulation during the testing period. Using a syringe mounted with a plastic tubing, a single drop of acetone was applied to the glabrous skin of the hindpaw of the animal once every 30 seconds, alternating between paws for a total of two applications per paw (4 total). The assay was digitally recorded and analyzed later by a blinded experimenter. To identify cool-induced nocifensive responses but not touch, the following scoring system was utilized: hind paw flinch was scored as a 1, as single lick was scored as a 2, multiple licks was scored as a 3, guarding, vocalization, and/or escape behaviors were scored as a 4. An average was then calculated across all four trials and used to represent a final score.

### Dry Ice Assay

Animals were placed in an elevated chamber with mesh flooring for all ‘habituation’ and testing periods. Following three ‘habituation’ sessions (20 minutes per day) in the behavior testing apparatus, a compacted pellet of dry ice was applied to the hindpaw of the animal once every 30 seconds, alternating between paws for a total of two applications per paw (4 total). The assay was digitally recorded and later analyzed by a blinded experimenter. To identify noxious cold nocifensive responses, a score was given one a scale of 0 to 2, where hindpaw flinch was scored as a 1, and one or more licks was scored as a 2. An average was then calculated across all four trials and used to represent a final score.

### Rectal Temperature Measurement

Briefly, the thumb and index fingers were used to gently grasp the nape of the neck, restraining the mouse for the duration of the testing period (1-2 minutes at most). A rectal thermometer (Right Temp Jr. Kent Scientific Corporation, Torrington, CT) was placed approximately 1.5 cm into the rectum, then held in place until a steady temperature could be recorded. For Calb1^Silenced^ experiments, the mouse was first weighed to determine the correct dosage of CNO (5 mg/kg), then the baseline rectal temperature was recorded, immediately followed by intraperitoneal injection of CNO, and then the final temperature was recorded 40 minutes later.

### C-Fos Induction

Following three ‘habituation’ sessions (20 minutes per day) in the behavior testing apparatus, a single acetone drop was applied to the hindpaw of *Calb1^Lbx1^*;Ai65 or *SOM^Cre^*;Ai14 mice once every 30 seconds over a 30-minute timespan. 1.5 hours later, mice were euthanized by isoflurane and perfused with 4% PFA in PBS. The lumbar spinal cord was then dissected, post-fixed for 2 hours at room temperature, cryoprotected for ≥ 24 hrs in sucrose solution (20% sucrose in PBS) at 4 °C, embedded in OCT, then sliced into 12-16 μm transverse sections on a cryostat (Leica Microsystems) for further c-Fos immunostaining (see *In Situ* Hybridization and Immunohistochemistry section above). For counting, the “peak c-Fos zone” was first identified, then a slice before and after was used for further quantification (ipsilateral and contralateral hemi-sections n = 9 each from 3 mice per experimental condition).

### Icilin Assay

As previously described (Dhaka et al., 2007), animals were habituated in a clear plexiglass experimental chamber for 20 minutes each day for two days prior to testing. On testing day, the mice were habituated for 20 minutes in the experimental chamber then 10 μl of 2.4 mg/ml icilin dissolved in 80% DMSO/20% PBS was injected into the right hindpaw. The mouse was then placed back into the behavior chamber and behavioral responses (hindpaw licking, flinching, and wet-dog shaking) were video-recorded for 60 minutes post-injection.

### Brainbow Virus Tracing

Schematic Image was created with BioRender.com (Figure 5B). Brainbow 3.0 AAV-PHP.eB was obtained from University of Michigan vector core. To systemically label dorsal spinal cord neurons, 50 μl of mCherry-TFP (1E12 gc total) was injected into the retro-orbital sinus of 3 mice. A new, clean needle will be inserted, bevel down, at an angle of approximately 45° through the inferior fornix conjunctival membrane (6 o’clock position into the eye socket). The needle was positioned behind the globe of the eye in the retro-bulbar sinus. After virus injection, the needle was gently removed to avoid injury to the eye. Six weeks after injection, adult mice were perfused with 1x PBS followed by 4% PFA. The spinal cords were dissected out and post-fixed in the 4% PFA overnight then used for expansion microscopy.

### Expansion Microscopy (ExM) and Immunohistochemistry

Fixed spinal cord ExM specimens were generated following the MiriEx expansion protocol (Shen et al., 2020). Briefly, adult mice were perfused with 1x PBS followed by 4% PFA. Spinal cords were dissected out, post-fixed in the 4% PFA overnight, then embedded in 2% low-melting agarose (Lanza, #50115) and vibratome (VT1200S, Leica, Germany) sectioned at 100μm. Representative sections were treated in 1.0 mM acrylic acid N-hydroxysuccinimide ester (AAx, Sigma, A8060) at 4 °C overnight, followed by 1x TBS washes for 3-4 hours (1 hour per wash). Subsequently the specimens were incubated in the MiriEx monomer solution (containing 5.3% Sodium Acrylate, 4% Acrylamide, 0.1% Bis acrylamide, 0.5% VA-44, and Triton X-100) at 4 °C overnight. The next morning, a 0.20mm iSpace (Sunjin Lab, IS312) was sealed onto a glass slide to create a gel chamber. The chamber was first half-filled with fresh 1x MiriEx monomer solution, next the tissues were gently laid flat in the chamber, then sealed with a glass cover slip and placed in a humidified container at 37 °C to allow gel polymerization for 2.5 hours. After polymerization, a clean razor blade was used to carefully trim away excess gel and scrape the gelled tissue off into a 2 ml centrifuge tube filled with denaturing buffer (200 mM SDS in 1x TBST) to allow protein denaturation at 70 °C overnight. The denatured tissues were washed with 0.1% PBST at least 4-5 times (1 hour per wash) at 50 °C. The washed tissue were then moved into new 1.5 ml centrifuge tubes for primary antibody staining (usually diluted 1:100 – 1:500) with: chicken anti-Homer (160006, Synaptic Systems, Goettngen, Germany), guinea pig anti-Bassoon (#141004, Synaptic Systems, Goettngen, Germany), rabbit anti-mCherry (generated by Cai lab, University of Michigan, Ann Arbor, MI, USA), rabbit anti-TFP (generated by Cai lab, University of Michigan, Ann Arbor, MI, USA), sheep anti-GFP (#4745-1051, Bio-Rad Laboratories, Hercules, CA, USA), then secondary antibody staining (1:1000) donkey anti-sheep Alexa fluor plus 488 (#713-546-147, Jackson Immunoresearch, West Grove, PA, USA), donkey anti-rabbit Alexa fluor Cy3 (#711-166-152, Jackson Immunoresearch, West Grove, PA, USA), donkey anti-guinea pig Alexa fluor 594 (#706-585-148, Jackson Immunoresearch, West Grove, PA, USA), donkey anti-chicken Alexa fluor 647 (#703-606-155, Jackson Immunoresearch, West Grove, PA, USA), followed by confocal imaging. A test round was performed to determine the number of staining rounds needed. If the morphology of the neurons could be completely defined by a single Brainbow fluorophore, then a single round of imaging was performed (synapse markers, Brainbow, and GFP). If the dendritic arbors partially overlapped, then two rounds of staining were performed (1: synaptic markers, GFP; 2: Brainbow markers, GFP; GFP channel was used for registration). For multi-round immunostaining and imaging, primary and secondary antibodies were stripped as previously described (Shen et al. 2020).

### Confocal Microscopy and Image Processing

The samples were gradually expanded 3-4 fold through multiple washes in 0.01x PBS. The expanded tissues were mounted in Poly-L-Lysine coated 6mm dishes (Corning, #356517). Confocal images were acquired with Zeiss LSM780 using a 20x 1.0 NA water immersion objective (421452-9800-000). The 32-channel GaAsP array detector was used to allow multi-track detection of four fluorophores with proper channel collection. All images were corrected for chromatic aberration using 0.5 mm TretraSpek fluorescent bead calibration images and the Detection of Molecules (DoM) ImageJ/Fiji plug in. Histogram matching was done to normalize intensity between z-slices in image stacks using the nTracer Align-Master ImageJ/Fiji plugin. nTracer, an ImageJ/Fiji plugin, was used to trace somas, dendrites, and axons of Brainbow labeled neurons (manual and tutorial videos can be found at https://www.cai-lab.org/ntracer-tutorial).

### Image Registration and Alignment Between Rounds

For experiments in which multiple rounds of immunostaining, imaging, and stripping were performed, image registration and alignment were implemented as previously described (Shen et al., 2020). Briefly, each round’s fiducial marker channel (GFP) was loaded into ImageJ/Fiji Big Warp plugin for rough, initial alignment. Elastix was then used to register the fiducial marker channels using B-splines. The resulting transformation was applied to each individual channel to create a merged image hyperstack. To register and align images from different rounds that are different expansion sizes, the lower resolution fiduciary channel was upsampled using bilinear interpolation to match the voxel size of the higher resolution fiduciary channel.

### Rabies Virus Tracing

As described previously (Pan et al., 2019), adult *Calb1::Cre;Lbx1^Flpo^, Rosa26^ds-HTB^* mice (n = 3) were anesthetized by isoflurane and a laminectomy was performed at the lumbar spinal cord (L3-L4). A fine needle was used to remove the dura matter and expose the spinal cord, then a fine glass capillary held by a nanoliter injector (WPI, Sarasota, FL) with stereotaxic device (David Kopf Instruments, Tujunga, CA) was inserted into the right side of the dorsal spinal cords. Focal injections of EnvA-pseudotyped, G-deleted-mCherry rabies virus (300 nl; ∼1E9 unit per ml, gift from Dr. Zhigang He at Boston Children’s Hospital) were injected into the dorsal spinal cord to target Calb1^+^ neurons in laminae I-III (50-300 μm depth from the surface) under the control of a micro-controller (Micro4, Sarasota, FL). Mice were perfused 10 days later and processed for further RNAscope experiments.

### Electrophysiology

#### Spinal Cord Slice Preparation

As described previously (Pan et al., 2019), the parasagittal spinal cord slices attached with the full length of the dorsal root and DRG were collected. Mice were deeply anesthetized with isoflurane, decapitated and the lumbar spinal cord was rapidly removed and placed in an ice-cold modified artificial cerebrospinal fluid (ACSF) containing (in mM): 80 NaCl, 2.5 KCl, 1.25, NaH_2_PO_4_, 0.5 CaCl_2_, 3.5 MgCl_2_, 25 NaHCO_3_, 75 sucrose, 1.3 sodium ascorbate and 3.0 sodium pyruvate, with pH at 7.4 and osmolality at 310-320 mOsm, bubbled with 95% O_2_ and 5% of CO_2_. Spinal cord slices (350-480 μm) attached with dorsal roots and DRGs were cut sagittally by a vibratome (VT1200S, Leica, Germany), as illustrated in Figure 7A. The slice was then incubated for about 1 hour at 33 °C in oxygenated (95% O_2_ and 5% CO_2_) cutting solution which contains (in mM): 125 NaCl, 2.5 KCl, 2 CaCl_2_, 1 MgCl_2_, 1.25 NaH_2_PO_4_, 26 NaHCO_3_, 25 D-glucose, 1.3 sodium ascorbate and 3.0 sodium pyruvate, with pH at 7.2 and osmolality at 310-320 mOsm. Icilin (1 μM), strychnine (2 μM), bicuculline (10 μM) and AMTB (100 μM) were diluted with a normal bath solution. All chemicals were purchased from MilliporeSigma (St. Louis, MO).

#### Whole-cell Patch Clamp Recordings

After incubation, spinal cord slices were placed in a recording chamber and perfused with oxygenated recording solution at a rate of 5 ml/minute at room temperature. Whole-cell recording experiments were then performed on Calb1^Lbx1^ dorsal horn neurons. Borosilicate glass pipettes (Sutter instrument, Novato, CA) with resistance of 3-6 MΩ were then filled with internal solution that contains (in mM): 130 potassium gluconate, 5 KCl, 4 Na_2_ATP, 0.5 NaGTP, 20 HEPES, 0.5 EGTA, pH 7.3 with KOH, and measured osmolality at 310-320 mOsm. Data were acquired by pClamp 10.0 software (Molecular Devices, San Jose, CA) with MultiClamp 700B patch clamp amplifier and Digidata 1550B (Molecular Devices, San Jose, CA). Responses were low pass filtered online at 2 kHz and digitized at 5 kHz.

#### Dorsal Root Stimulation

Different responses of dorsal horn neurons to primary afferent inputs were recorded under different recording conditions. Firstly, evoked excitatory postsynaptic currents (eEPSCs) were detected by holding membrane potential at −70 mV, which minimized evoked inhibitory postsynaptic currents (eIPSCs) (Yoshimura and Jessell, 1990). Whether a neuron receives Aβ, Aδ or C-fiber inputs directly (mono-eEPSC) or indirectly (poly-eEPSC) were determined under this recording condition. Monosynaptic inputs for Aβ, Aδ or C fibers were determined by high frequency stimulation at 20, 2 or 1 Hz, respectively (Pan et al., 2019; Torsney and MacDermott, 2006). Transduction velocity was also used to determine monosynaptic inputs: Aβ, 2.16-4.06 m/s; Aδ, 0.92-1.04 m/s; C, 0.18-0.62 m/s. Secondly, eIPSCs were recorded by holding membrane potential at 0mV when eEPSCs were minimized. Bicuculline (10 µM, MilliporeSigma, St. Louis, MO) and/or strychnine (2 μM, MilliporeSigma, St. Louis, MO) were used to disinhibit dorsal horn neurons. Thirdly, dorsal root stimulation-evoked IPSP, EPSP, or APs were detected by current clamp recording at the resting membrane potential. Action potential firing patterns were determined by current clamp recording at the resting membrane potential.

#### Characterization of Firing Pattern

The steady-state firing pattern was isolated from the initial transient phase, i.e. the firing pattern immediately after the beginning of the current phase. There were three main starting patterns characterized: The onset was indistinguishable from the rest of the spike response (tonic), neurons responded with a much greater frequency of spikes in the transient (initial burst) than in the steady state, and neuron firing started with a delay (delay). After an initial transient, neurons displayed a steady-state pattern. Again, there were three main types: regularly spaced spikes (tonic), gradually increasing interspike interval (adapting), or regular alternating between short and long intervals (bursting).

### Single Cell RT-PCR

PCR-amplified cDNA libraries for single cells were generated from individual spinal cord neuron cells (SuperScript™ IV Single Cell cDNA PreAmp, Cat: 11752048, Thermo, CA). The cDNA quality of each cell was confirmed by PCR for GAPDH and Somatostatin (SOM). Primers were designed with primer-BLAST of the GAPDH and SOM genes (Table 1). 1x reaction mix, 2 mM MgCl_2_, 250 μM each deoxynucleotide for each reaction; (Thermo, CA), 0.25 μM forward primer, 0.25 μM reverse primer and 2.5U SuperScript™ One-Step RT-PCR (Thermo, CA) were combined with 1 μl template cDNA. PCRs for SOM and GAPDH were performed with 35 cycles of initial 10-minute denaturation (94 °C), 30-second denaturation (94 °C), 30-second annealing (55 °C), and 2-minute extension (72 °C). After 10-minute post-elongation (72 °C). Table 1 shows the annealing temperature range of 35-cycle (SOM, GAPDH), the annealing temperature range of 20-cycle. Amplified products were run on 1.5% agarose gels. Certain bands were observed in the control spinal cord neurons, but no bands were seen in the water (AM9935, Thermo, CA).

**Table 1.**
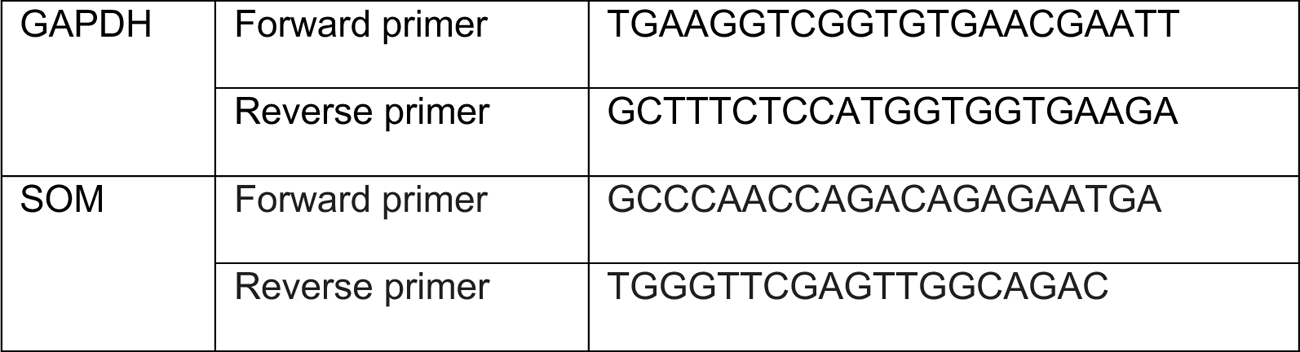
Primers for RT-PCR

### Ca^2+^ Imaging

As previously described (Pan et al., 2019), acutely dissociated DRG neurons from 4 mice were first incubated in at 37 °C for 30 minutes in 4 mm Fura-2-acetoxymethyl ester and 0.2% Pluronic® F127 (Thermo Fisher Scientific, Waltham, MA), washed three times, then incubated for 20 minutes in standard extracellular solution (in mM): 10 HEPES [pH 7.4], 5 KCl, 140 NaCl, 2 MgCl_2_, 2 CaCl_2_, and 10 glucose. Calcium imaging was performed using a 20x water-immersion objective with high transmission efficiency of UV light (Olympus IX73, Japan). A Roper Cool-Snap CCD camera was used for image acquisition, and images were processed with MetaFluor software (Molecular Devices, San Jose, CA). Icilin (in μM): 0.1, 0.3, 1, 3, 5, and 10, and KCl (50 mM) were dissolved in standard extracellular solution. Temperature was maintained and verified using Heating and Cooling Application Fundamentals (Warner Instruments LLC, Hamden, CT).

### Tracing SPB Neurons

Adult *Calb1^Lbx1^;*Ai65 mice were anesthetized by isoflurane and a craniotomy was performed. To mark SPB neurons, we bilaterally injected cholera-toxin B (2%, #C34775, #C34778, ThermoFisher, Waltham, MA), a retrograde tracer, into the lateral parabrachial nucleus (lPB) in *Calb1^Lbx1^;*Ai65 mice. A nanoliter injector (WPI, Sarasota, FL) coordinated with a stereotaxic device (David Kopf Instruments, Tujunga, CA) was positioned at the coordinates and a dental drill (8149285, Meisingerusa, Centennial, CO) was used to expose the brain. Next, a fine glass capillary containing CTB, was inserted bilaterally into the lPB and a microcontroller (micro4, Sarasota, FL) was used to deliver 1-1.5 μl of CTB to each region. To label Calb1^Lbx1^-Tomato cooling-sensitive SPB neurons, 6-10 days after surgery, 3 *Calb1^Lbx1^;*Ai65 mice were habituated to the behavior chamber for two days before acetone-induced c-Fos protocol (as previously described). For electrophysiological recordings of Calb1^Lbx1^-Tomato SPB neurons, 5 control and 4 Calb1^Abl^ mice were injected in the lPB with CTB, then 7–14 days later the spinal cord and DRG were dissected out together and recorded.

### Quantification and Statistical Analysis

Results are expressed as mean ± SEM. Statistical analysis was performed in Prism 6 or 9 (GraphPad). A threshold of p < 0.05 was accepted as statistically different and p > 0.05 considered non-significant. For ablation experiments, locomotion coordination, touch, acute pain assessment, and temperature (except two temperature preference and gradient temperature) data were subjected to Student’s t tests. Temperature preference assay and gradient temperature data were subjected to two-way ANOVA with Sidak and Bonferroni post hoc analyses respectively. For silencing experiments, all behavior data were subjected to two-way ANOVA with Sidak post hoc analysis. C-Fos data were subjected to two-way ANOVA with Sidak post hoc analysis. For electrophysiological results, data were analyzed with two-way ANOVA with Tukey (AMTB application; percent Icilin Responding cells; percent icilin-induced APs in icilin responding cells) and Sidak (Current by icilin) post hoc analyses. No statistical methods were used to predetermine sample sizes, but our sample sizes are similar to those reported in previous publications (Duan et al., 2014; Pan et al., 2019). Sex differences were analyzed (two-way ANOVA with Sidak post hoc analysis) and no statistical significance was determined (data not shown).

## Data and Code Availability

The MATLAB script for the temperature gradient analysis is available upon request.

