## Supplementary Figures for "A Spinal Circuit That Transmits Innocuous Cool Sensations"

1. Department of Molecular, Cellular, and Developmental Biology, University of Michigan, Ann Arbor, MI 48109, USA.
2. Neuroscience Graduate Program, University of Michigan, Ann Arbor, MI, 48019, USA
3. Department of Cellular Developmental Biology, University of Michigan, Ann Arbor, MI 48109, USA.
4. Life Sciences Institute, University of Michigan, Ann Arbor, MI 48109, USA; Department of Molecular and Integrative Physiology, University of Michigan, MI 48109, USA.
5. Department of Mechanical Engineering, University of Michigan, Ann Arbor, MI 48109, USA.
6. Department of Electrical Engineering and Computer Science, University of Michigan, Ann Arbor, MI 48109, USA.
7. These authors contributed equally.

#### SUPPLEMENTARY FIGURE LEGENDS

**Figure S1. Classification of NK1R$^+$ Neuronal Quantification (related to Figure 1).**

Triple staining of Calb1$^{Lbx1}$-Tomato$^+$ signals (red) with NK1R immunostaining (green). Colocalization with DAPI (blue) was used to identify NK1R$^+$ projection neurons. Insets represent higher magnification of the boxed area. Arrowhead indicates NK1R and DAPI double-labeled cell, but not Tomato$^+$. n = 18 sections. Scale bar, 100 $\mu$m.

**Figure S2. Ablation of Calb1$^{Lbx1}$ Neurons in the Spinal Cord Does Not Affect the Expression of Calb1$^+$ Neurons in the Brain (related to Figure 2).**

**(A)** Schematic demonstrating intersectional genetic ablation strategy to express diphtheria toxin receptor in Calb1$^{Lbx1}$ neurons to selectively ablate the Calb1$^{Lbx1}$ population in adulthood.

**(B)** Ablation of Calb1$^{Lbx1}$ neurons does not affect the expression of Calb1$^{Cre}$-Tomato$^+$ neurons in the parabrachial nucleus (top panels) or the somatosensory cortex region 1 (S1) (bottom panels) in control and Calb1$^{Abl}$ mice. Scale bar: 100 $\mu$m.

**Figure S3. Behavioral Assessment of Locomotion, Touch, and Nociceptive Sensations in Calb1$^{Abl}$ Mice (related to Figure 2).**

**(A)** Bar graph represents latency to fall in control and Calb1$^{Abl}$ animals in the Rotarod assay. No significant difference in the Rotarod assay. Control: n = 15; Calb1$^{Abl}$: n = 16; ns, no significant difference; Student's unpaired t test.

**(B)** Light-brushing evoked a significantly higher score in control compared to Calb1$^{Abl}$ mice (Control: n = 10; Calb1$^{Abl}$: n = 9; * $p < 0.05$; Mann-Whitney test).

**(C)** No significant difference in latency to respond to sticky tape administration in control and Calb1$^{Abl}$ mice was observed. Control: n = 7; Calb1$^{Abl}$: n = 8; ns, no significance; Student's unpaired t test.

**(D)** Latency to respond to pinch was not significantly different between control and Calb1$^{Abl}$ mice. n = 21 in each group; ns, no significance; Student's unpaired t test.

**(E)** Pinprick assay was not significantly different in control compared to Calb1$^{Abl}$ mice. Control: n = 13; Calb1$^{Abl}$: n = 8; ns, no significance; Student's unpaired t test.

**(F)** Acute punctate mechanical pain as measured by von Frey response rate threshold across increasing force. Control: n = 10; Calb1$^{Abl}$: n = 9; *** $p < 0.001$, **** $p < 0.0001$, two-way ANOVA with Sidak post hoc analysis.

**(G-H)** Noxious heat thermosensation was measured through Hargreaves assay **(G)** and hot plate assay set to 46 °C, 50 °C, or 54 °C **(H)**. No significant difference in latency to flinch the front paw or lick the hindpaw between control and Calb1$^{Abl}$ animals. n = 6-13 in each group; ns, no significance, two-way ANOVA with Sidak post hoc analysis.

**Figure S4. Behavioral Assessment of Locomotion, Touch, and Nociceptive Sensations in Calb1$^{Silenced}$ Mice (related to Figure 3).**

**(A)** Schematic showing the intersectional genetic strategy to temporally restrict the silencing of spinal Calb1$^{Lbx1}$ neurons.

**(B)** Locomotor agility as measured by latency to fall in the rotarod assay was not significantly different between control and Calb1$^{Silenced}$ mice, or between Calb1$^{Silenced}$ mice before compared to 40 minutes after CNO administration. Control: n = 11; Calb1$^{Silenced}$: n = 9; ns, no significant differences, two-way ANOVA with Sidak post hoc analysis.

**(C)** Calb1$^{Silenced}$ mice displayed a significant deficit in light brushing-evoked responses 40 minutes after CNO administration within group (Calb1$^{Silenced}$ before CNO administration) and compared to control animals. Control: n = 11; Calb1$^{Silenced}$: n = 9; **** $p < 0.0001$, two-way ANOVA with Sidak post hoc analysis.

**(D)** Latency to respond to sticky tape administration was not significantly altered 40 minutes after CNO administration in Calb1$^{Silenced}$ mice compared to baseline (before CNO administration) and control animals. Control: n = 11; Calb1$^{Silenced}$: n = 9; ns, no significant difference, two-way ANOVA with Sidak post hoc analysis.

**(E)** No significant difference in pinch response latency was observed in Calb1$^{Silenced}$ mice compared to before CNO administration or controls animals. Control: n = 11; Calb1$^{Silenced}$: n = 9; ns, no significant difference, two-way ANOVA with Sidak post hoc analysis.

**(F)** Pinprick assay was not significantly different in Calb1$^{Silenced}$ mice compared to littermate control or across time (before compared to 40 minutes after CNO administration). Control: n = 11; Calb1$^{Silenced}$: n = 9; ns, no significant difference, two-way ANOVA with Sidak post hoc analysis.

**(G-H)** Noxious heat thermosensation was measured through Hargreaves assay **(G)** and hot plate assay at 46 °C, 50 °C, or 54 °C **(H)**. There was no significant difference in the latency to flinch the front paw of lick the hindpaw between Calb1$^{Silenced}$ mice across time before compared to 40 minutes after CNO administration or compared to littermate controls. Control: n = 11; Calb1$^{Silenced}$: n = 9; ns, no significant difference, two-way ANOVA with Sidak post hoc analysis.

**Figure S5. SOM$^{Cre}$-Tomato neurons are not cooling-sensitive (related to Figure 4).**

**(A)** Double staining of c-Fos and Tomato$^{+}$ signals in the ipsilateral (left) and contralateral (right) dorsal horn of acetone-treated *SOM$^{Cre}$*;Ai14 mice. Inset (middle) represents higher magnification of the boxed area. Arrowheads shows a cell positive for Tomato alone. Scale bar, 100 μm. Very little overlap was observed. n = 9 from each hemi-section.

**(B)** Quantified forelimb flinch and lick withdrawal latency to 0 °C cold plate. Control: n = 10; SOM$^{Abl}$: n = 9; ns, no significant difference; two-way ANOVA with Sidak post hoc analysis.

**Figure S6. Calb1$^{Cre}$ neurons in the superficial dorsal horn receive monosynaptic inputs from TRPM8$^{+}$ primary sensory neurons (related to Figure 5).**

**(A)** Quantification of nocifensive responses (lick or scratch) after icilin injection into the plantar region of the hindpaw in control and Calb1$^{Abl}$ mice. control: n = 7; Calb1$^{Abl}$: n = 9; no significance, two-way ANOVA with Sidak post hoc analysis.

**(B)** Retro-orbital injection of *Cre*-dependent AAV-PHP.eB Brainbow in *Calb1$^{Cre}$; TRPM8$^{GFP}$* mice to selectively infect the central nervous system and not the periphery (DRG). TRPM8$^{GFP}$: Green; mCh: Brainbow expressing mCherry; TFP: Brainbow expressing Teal Fluorescent Protein.

**(C)** TRPM8$^{+}$ presynaptic neurons formed multiple synaptic connections in close proximity with Calb1$^{Brainbow}$ postsynaptic spinal neurons. Arrowhead indicates quadruple positive synaptic pairs (red: presynaptic marker Bassoon; green: postsynaptic marker Homer; purple: presynaptic neuron TRPM8$^{+}$; blue: postsynaptic neuron Calb1$^{Brainbow}$). Arrowhead indicates an orphan synaptic pair (colocalized pre- and post- synaptic markers only).

**(D)** Schematic adapted from Pan et al., demonstrating strategy for monosynaptic labeling of pre-synaptic neurons after selective infection of Calb1$^{+}$ neurons (GFP$^{+}$) with pseudotyped EnvA-mCherry rabies virus (RV-mCherry$^{+}$) in the dorsal spinal cord in a HTB transgenic mouse line.

**(E)** Dorsal spinal cord after rabies virus infection of *Calb1$^{Lbx1}$;* HTB animals (HTB-GFP: green). Arrow indicates an infected Calb1$^{+}$ neuron (GFP$^{+}$ and mCherry$^{+}$). Arrowhead indicates a presynaptic neuron of Calb1$^{+}$ infected neurons (mCherry$^{+}$ only). Scale bar: 100 $\mu$m.

**(F)** Representative image of the DRG showing infected presynaptic neurons (Rabies virus labeled mCherry$^{+}$ denoted as RabV-mCh) and RNAscope-labeled *TRPM8* mRNA (green) neurons. Arrow indicates rabies virus infected Rabies virus labeled mCherry$^{+}$ (denoted as RabV-mCh) colocalized with a TRPM8$^{+}$ (green) presynaptic peripheral sensory neurons. Scale bar: 100 $\mu$m.

**Figure S7. Activation of superficial Calb1$^{Lbx1}$ neurons after icilin application to the DRG is mediated by C-fiber stimulation while Calb1$^{Lbx1}$;SOM$^{+}$ neurons receive Aβ, Aδ, and C fiber inputs (related to Figure 7).**

**(A)** Exemplar trace of calcium imaging change in fluorescence responses (F340/380) to various stimuli in cultured TRPM8$^{GFP}$ DRG neurons. KCl: Potassium Chloride (50 mM). Icilin concentrations (μM): 0.1, 0.3, 1, 3, 5, and 10.

**(B)** Quantification of the percent of cells that respond to each stimulus in TRPM8$^{GFP}$ cultured DRG neurons. TRPM8$^{GFP}$ neurons: n = 103; icilin responding neurons: n = 84; **** $p < 0.0001$, two-way ANOVA with Tukey post hoc analysis. KCl: Potassium Chloride. Icilin concentrations (μM): 0.1, 0.3, 1, 3, 5, and 10.

**(C)** The total percentage of all DRG neurons that were responsive at various stimuli. DRG neurons: n = 697; icilin-responding neurons: n = 84; **** $p < 0.0001$, two-way ANOVA with Tukey post hoc analysis). KCl: Potassium Chloride. Icilin concentrations (μM): 0.1, 0.3, 1, 3, 5, and 10.

**(D)** Schematic demonstrating two-chamber apparatus that contains a DRG chamber and spinal cord chamber. Neurons are recorded in the spinal cord chamber while the attached DRG is separately housed to isolate stimulation application (electrical, icilin, and/or AMTB) (top). Glycine and $GABA_A$ receptor antagonists, strychnine and bicuculine respectively were administered to the spinal cord chamber (bottom) to block any inhibitory gating. Calb1$^{Lbx1}$;SOM$^{-}$ neurons were recorded in the superficial lamina I-$II_o$ whereas Calb1$^{Lbx1}$;SOM$^{+}$ neurons were recorded from lamina II. Red dots represent recorded neurons.

**(E)** Calb1$^{Lbx1}$-Tomato$^{+}$ neurons recorded in lamina I-$II_o$ upon icilin administration to either the spinal cord chamber (top row) or DRG chamber (bottom row). A small icilin induced EPSCs (left column) and no APs (right column) responses were recorded from icilin application to the spinal cord chamber whereas icilin application to the DRG chamber evoked both EPSCs and APs. Blue line: 1 μM icilin application. n = 10.

**(F-H)** Exemplar traces of Aβ-evoked **(F)**, Aδ-evoked **(G)**, and C-evoked **(H)** EPSCs, IPSCs, APs recorded under normal conditions (top row) and under disinhibition condition (bottom row) in $Calb1^{Lbx1}$;$SOM^-$ neurons in lamina I-$II_o$. Str, strychnine. Bic, Bicuculline. Red arrowheads indicate stimulation artifacts.

**(I-K)** Exemplar traces of Aβ-evoked **(I)**, Aδ-evoked **(J)**, and C-evoked **(K)** EPSCs, IPSCs, APs recorded under normal conditions (top row) and under disinhibition condition (bottom row) in $Calb1^{Lbx1}$;$SOM^+$ neurons in lamina II. Str, strychnine. Bic, Bicuculline. Red arrowheads indicate stimulation artifacts.

**Figure S8. RT-PCR mediated identification of $Calb1^{Lbx1}$;$SOM^+$ and $Calb1^{Lbx1}$;$SOM^-$ neurons**

**(A)** Following whole-cell patch clamp recordings, all recorded $Calb1^{Lbx1}$;$SOM^-$ neurons in lamina I-$II_o$ were identified by RT-PCR. SOM: Somatostatin; GAPDH: reference gene. M, monosynaptic inputs; P, polysynaptic inputs; +, positive control; -, negative control (mastermix without template DNA). Red highlights these neurons generating both Icilin-EPSCs and APs. Data were collected from 10 mice.

**(B)** Following whole-cell patch clamp recordings, all recorded $Calb1^{Lbx1}$;$SOM^+$ neurons in lamina II were identified by RT-PCR. SOM: Somatostatin; GAPDH: reference gene. M, monosynaptic inputs; P, polysynaptic inputs; +, positive control; -, negative control (mastermix without template DNA). Data were collected from 10 mice.

### Figure S1

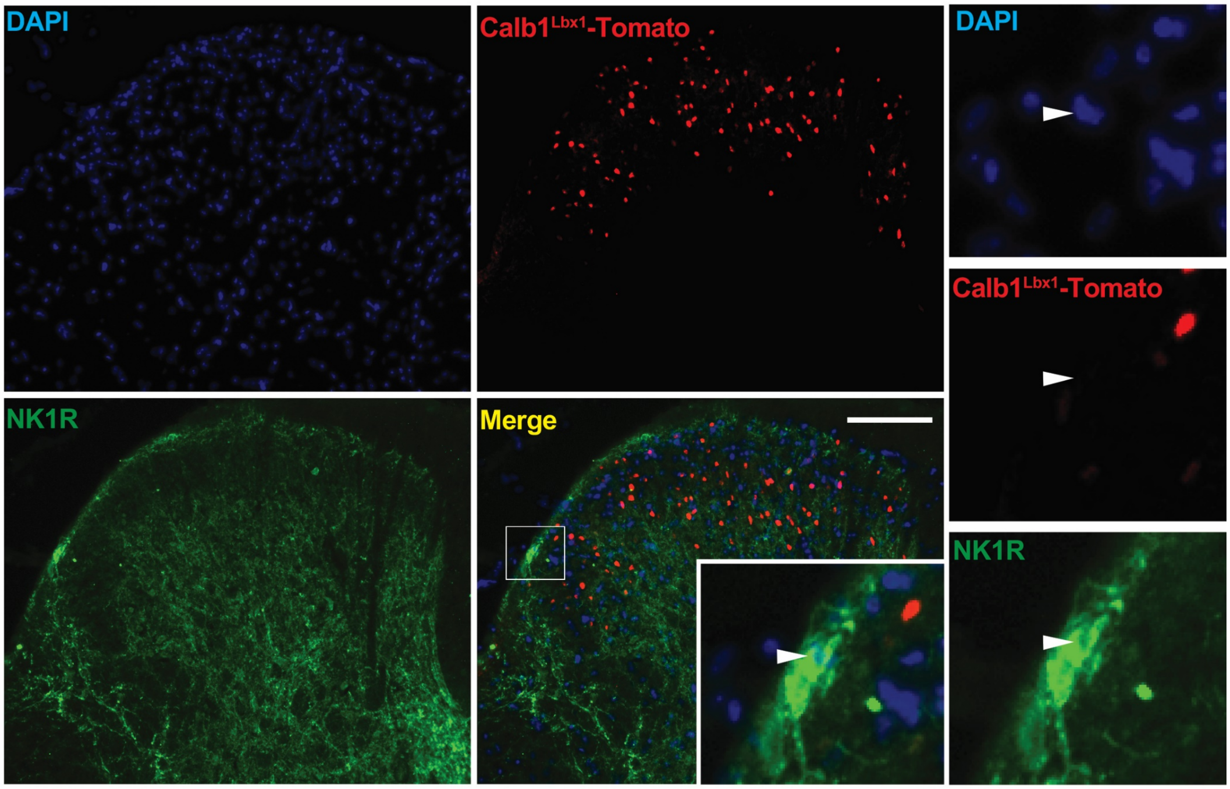
DAPI
Calb1^Lbx1-Tomato
DAPI
Calb1^Lbx1-Tomato
NK1R
Merge
NK1R

#### Figure S2

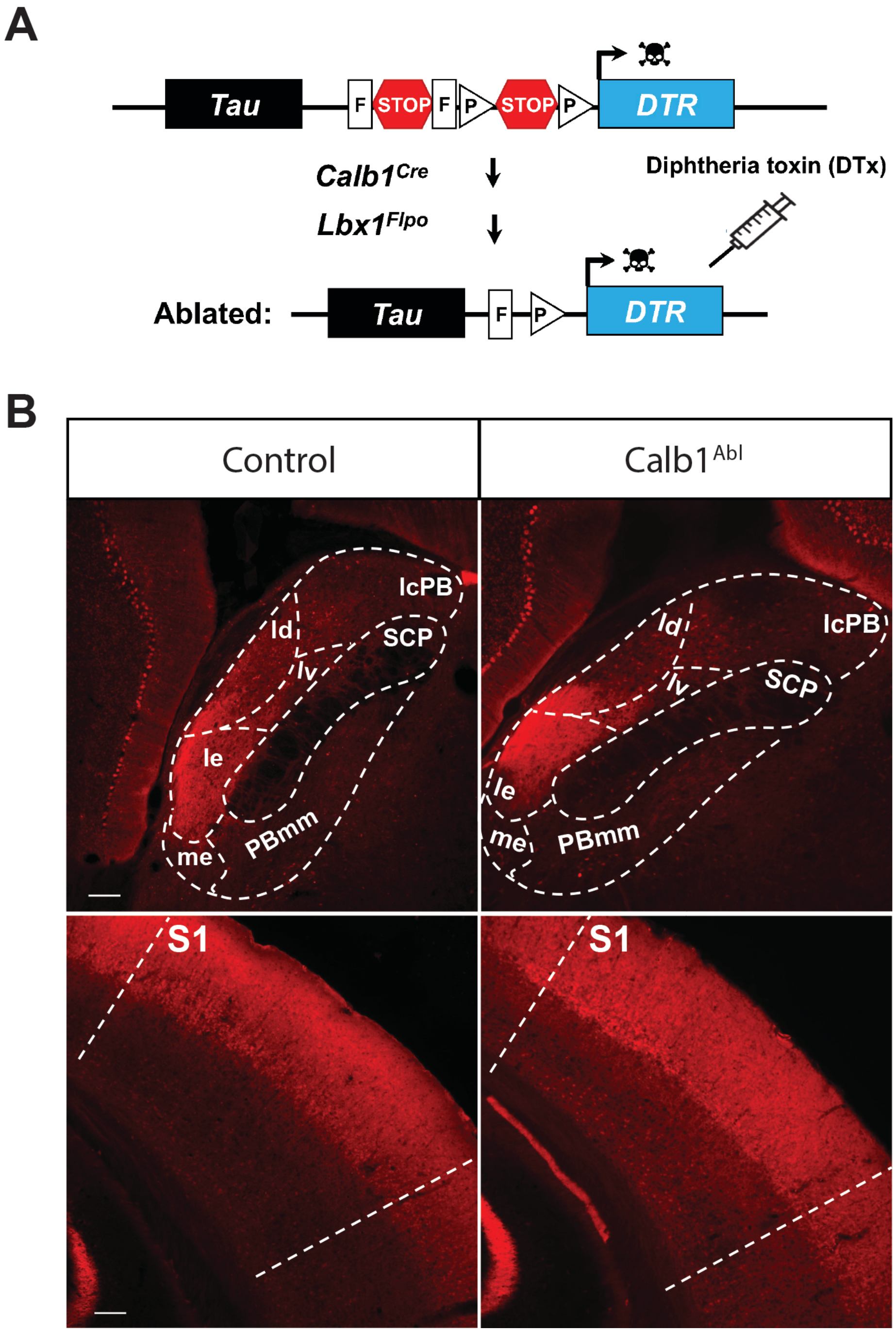
A
Tau
F
STOP
F
P
STOP
P
DTR
Calb1Cre
Lbx1Flpo
Diphtheria toxin (DTx)
Ablated:
Tau
F
P
DTR
B
Control
Calb1 Abl
lcPB
ld
SCP
lv
le
me
PBmm
lcPB
ld
lv
SCP
le
me
PBmm
S1
S1

#### Figure S3

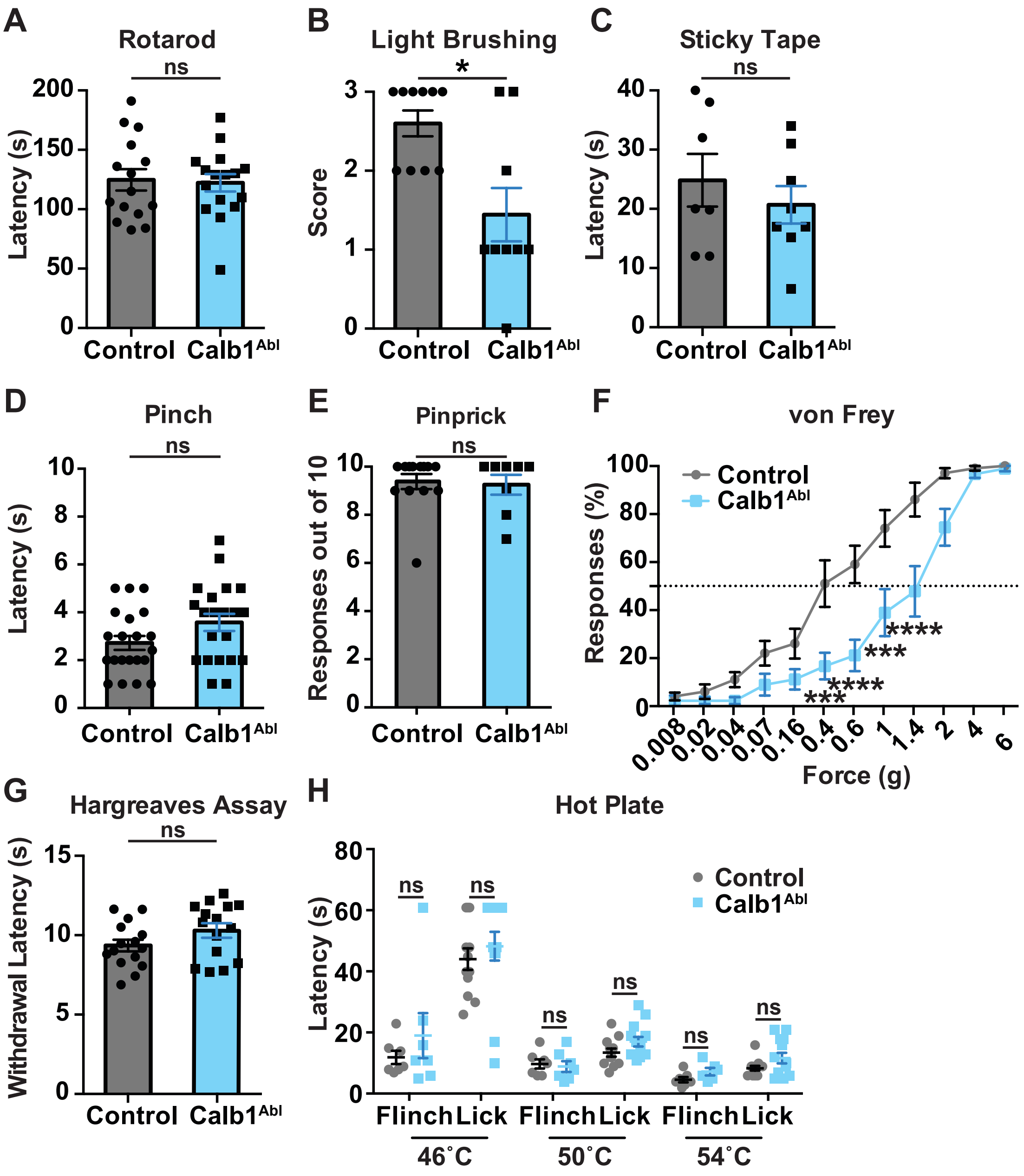
A
Rotarod
ns
Latency (s)
200
150
100
50
0
Control
Calb1Abl
B
Light Brushing
*
Score
3
2
1
0
Control
Calb1Abl
C
Sticky Tape
ns
Latency (s)
40
30
20
10
0
Control
Calb1Abl
D
Pinch
ns
Latency (s)
10
8
6
4
2
0
Control
Calb1Abl
E
Pinprick
ns
Responses out of 10
10
8
6
4
2
0
Control
Calb1Abl
F
von Frey
Control
Calb1Abl
Responses (%)
100
80
60
40
20
0
****
***
****
***
0.008
0.02
0.04
0.07
0.16
0.4
0.6
1
1.4
2
4
6
Force (g)
G
Hargreaves Assay
ns
Withdrawal Latency (s)
15
10
5
0
Control
Calb1Abl
H
Hot Plate
Control
Calb1Abl
Latency (s)
80
60
40
20
0
ns
ns
ns
ns
ns
ns
Flinch
Lick
Flinch
Lick
Flinch
Lick
46˚C
50˚C
54˚C

#### Figure S4

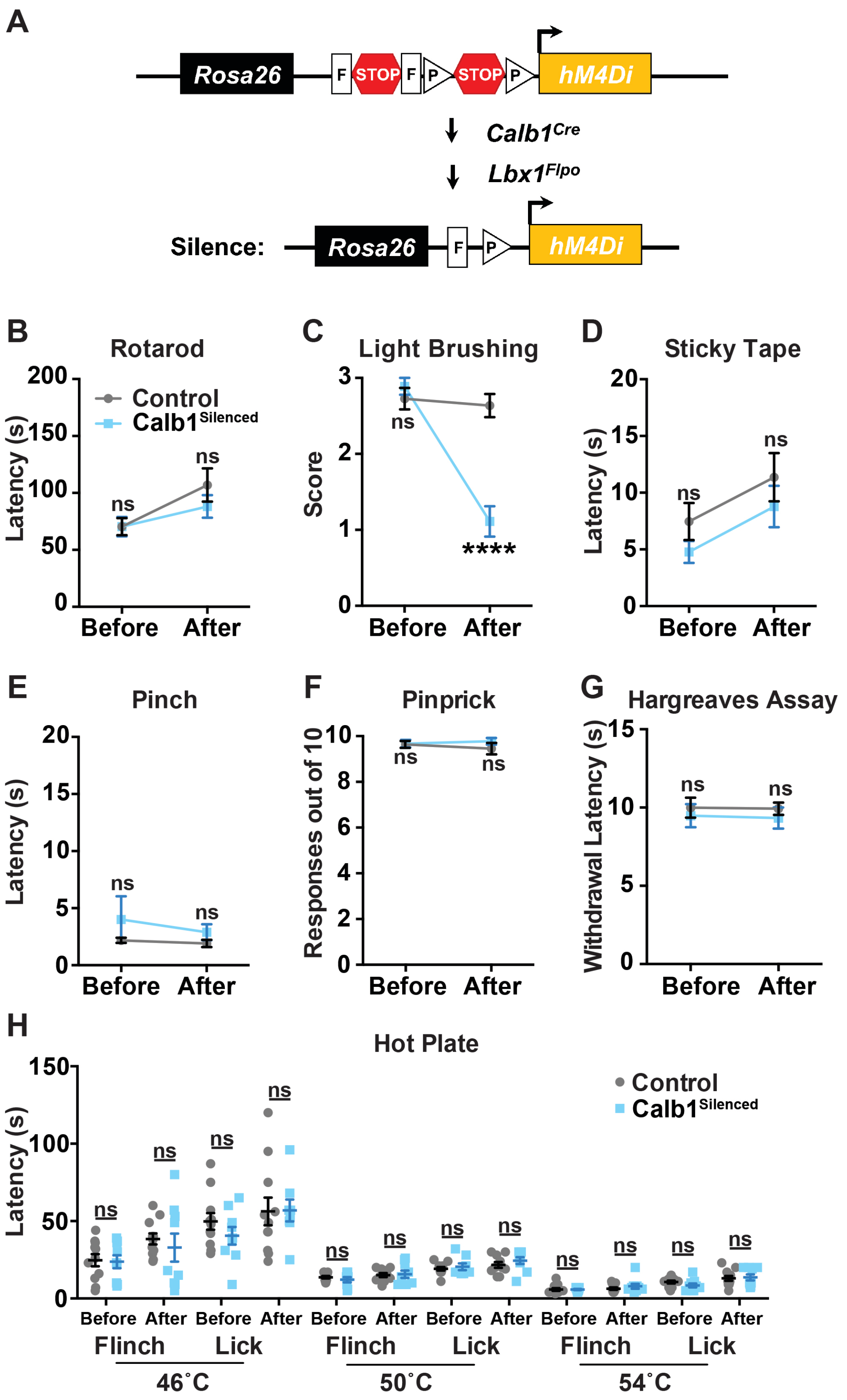
A
Rosa26
F
STOP
F
P
STOP
P
hM4Di
Calb1Cre
Lbx1Flpo
Silence:
Rosa26
F
P
hM4Di
B
Rotarod
Latency (s)
Control
Calb1Silenced
ns
ns
Before After
C
Light Brushing
Score
ns
****
Before After
D
Sticky Tape
Latency (s)
ns
ns
Before After
E
Pinch
Latency (s)
ns
ns
Before After
F
Pinprick
Responses out of 10
ns
ns
Before After
G
Hargreaves Assay
Withdrawal Latency (s)
ns
ns
Before After
H
Hot Plate
Latency (s)
Control
Calb1Silenced
ns
Before After Before After Before After Before After Before After Before After
Flinch Lick Flinch Lick Flinch Lick
46˚C
50˚C
54˚C

#### Figure S5

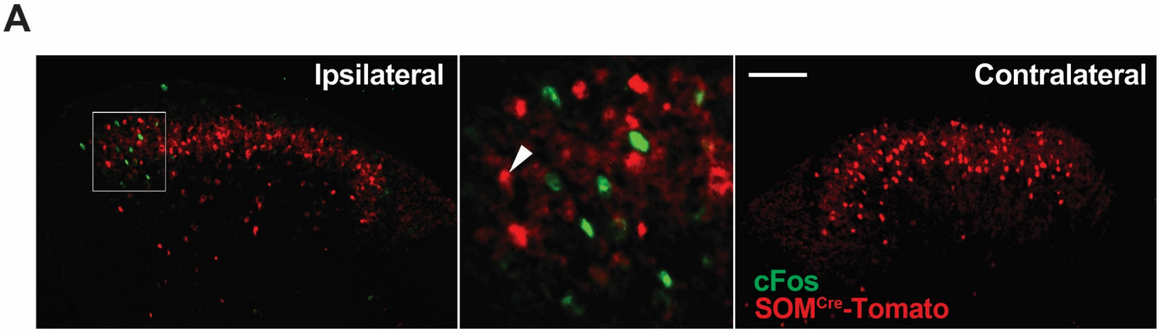

A
Ipsilateral
Contralateral
cFos
SOMCre-Tomato


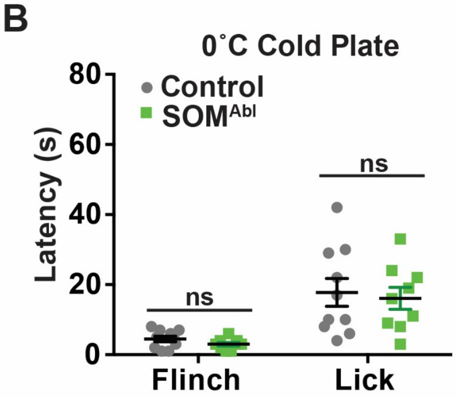

B
0˚C Cold Plate
Control
SOMAbl
Latency (s)
80
60
40
20
0
ns
ns
Flinch
Lick

#### Figure S6

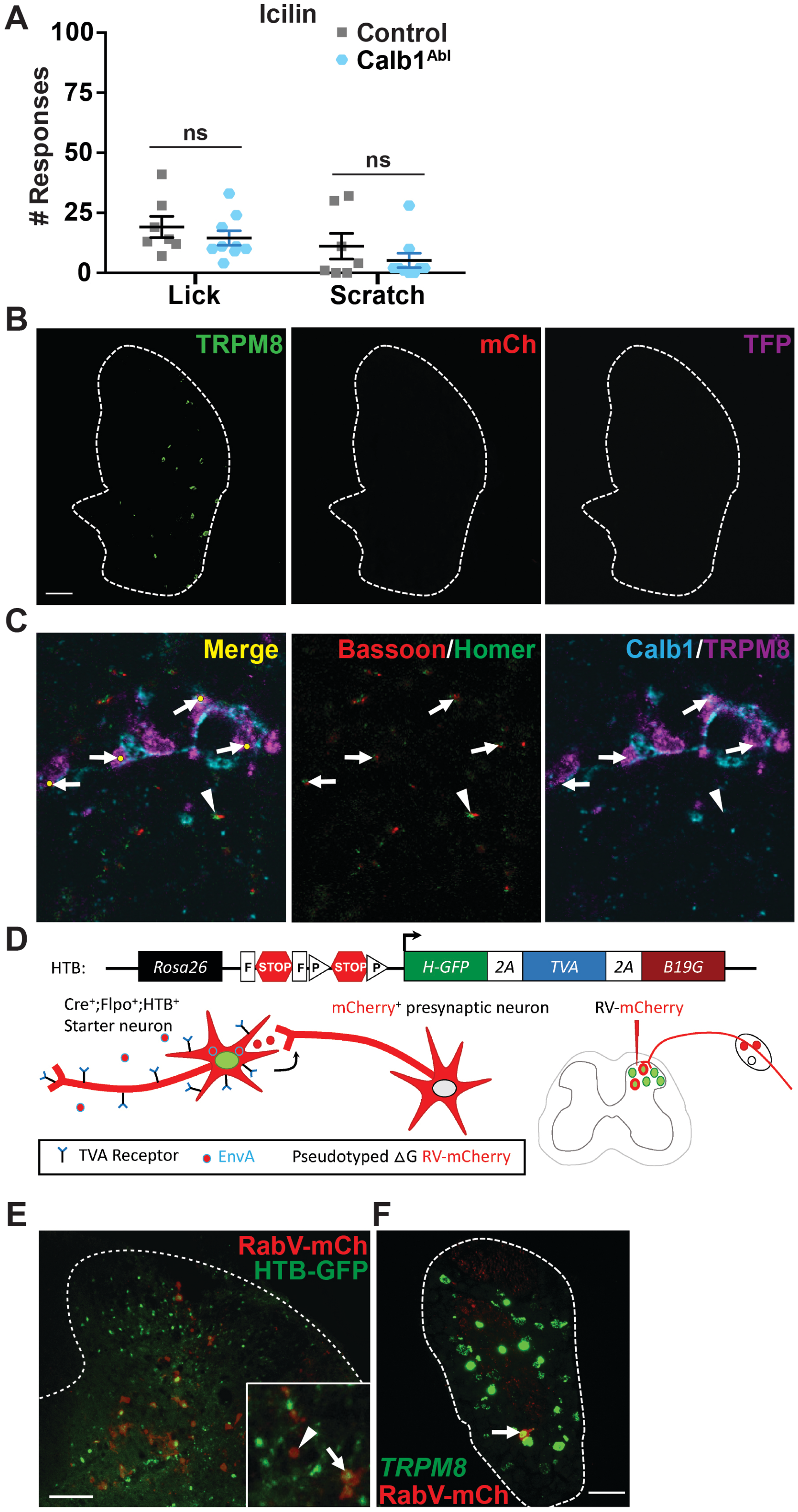
A
Icilin
Control
Calb1Abl
### Responses
100
75
50
25
0
ns
ns
Lick
Scratch
B
TRPM8
mCh
TFP
C
Merge
Bassoon/Homer
Calb1/TRPM8
D
HTB:
Rosa26
F
STOP
F
P
STOP
P
H-GFP
2A
TVA
2A
B19G
Cre+;Flpo+;HTB+
Starter neuron
mCherry+ presynaptic neuron
RV-mCherry
TVA Receptor
EnvA
Pseudotyped ΔG RV-mCherry
E
RabV-mCh
HTB-GFP
F
TRPM8
RabV-mCh

#### Figure S7

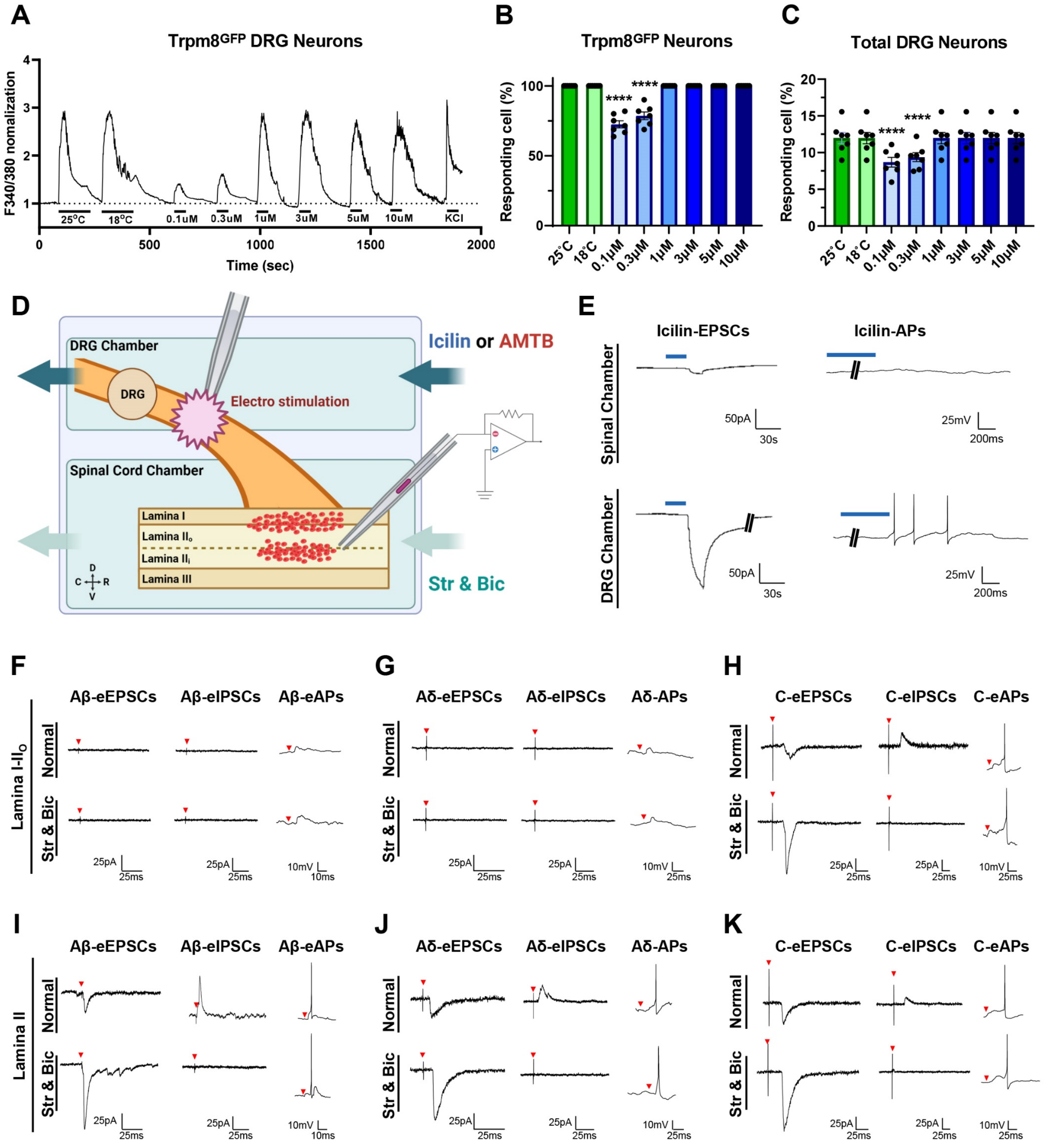
A
Trpm8GFP DRG Neurons
F340/380 nomalization
25°C
18°C
0.1uM
0.3uM
1uM
3uM
5uM
10uM
KCl
Time (sec)
B
Trpm8GFP Neurons
Responding cell (%)
C
Total DRG Neurons
Responding cell (%)
D
DRG Chamber
DRG
Electro stimulation
Icilin or AMTB
Spinal Cord Chamber
Lamina I
Lamina IIo
Lamina IIi
Lamina III
Str & Bic
E
Icilin-EPSCs
Icilin-APs
Spinal Chamber
DRG Chamber
F
Aβ-eEPSCs
Aβ-eIPSCs
Aβ-eAPs
Lamina I-IIo
Normal
Str & Bic
G
Aδ-eEPSCs
Aδ-eIPSCs
Aδ-APs
H
C-eEPSCs
C-eIPSCs
C-eAPs
I
Lamina II
J
K

#### Figure S8

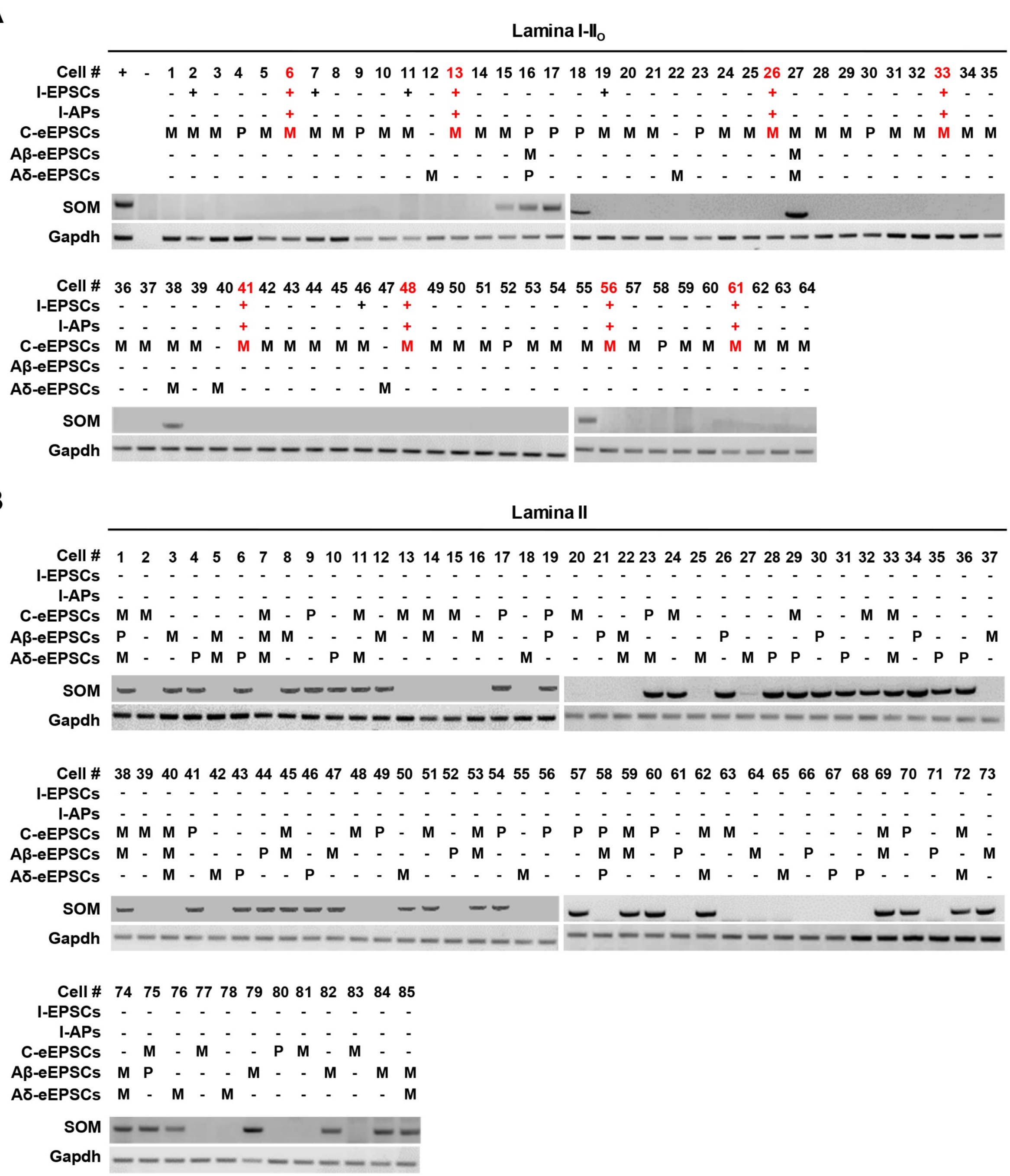

A
Lamina I-II$_O$
Cell # + - 1 2 3 4 5 6 7 8 9 10 11 12 13 14 15 16 17 18 19 20 21 22 23 24 25 26 27 28 29 30 31 32 33 34 35
I-EPSCs - + - - - + + - - - + - + - - - - - + - - - - - - + - - - - - - + - -
I-APs - - - - - + - - - - - - + - - - - - - - - - - - - + - - - - - - + - -
C-eEPSCs M M M P M M M M P M M - M M M P P P M M M - P M M M M M M P M M M M M
Aβ-eEPSCs - - - - - - - - - - - - - - - M - - - - - - - - - - M - - - - - - - -
Aδ-eEPSCs - - - - - - - - - - - M - - - P - - - - - M - - - - M - - - - - - - -
SOM
Gapdh
Cell # 36 37 38 39 40 41 42 43 44 45 46 47 48 49 50 51 52 53 54 55 56 57 58 59 60 61 62 63 64
I-EPSCs - - - - - + - - - - + - + - - - - - - - + - - - - + - - -
I-APs - - - - - + - - - - - - + - - - - - - - + - - - - + - - -
C-eEPSCs M M M M - M M M M M M - M M M M P M M M M M P M M M M M M
Aβ-eEPSCs - - - - - - - - - - - - - - - - - - - - - - - - - - - - -
Aδ-eEPSCs - - M - M - - - - - - M - - - - - - - - - - - - - - - - -
SOM
Gapdh
B
Lamina II
Cell # 1 2 3 4 5 6 7 8 9 10 11 12 13 14 15 16 17 18 19 20 21 22 23 24 25 26 27 28 29 30 31 32 33 34 35 36 37
I-EPSCs - - - - - - - - - - - - - - - - - - - - - - - - - - - - - - - - - - - - -
I-APs - - - - - - - - - - - - - - - - - - - - - - - - - - - - - - - - - - - - -
C-eEPSCs M M - - - - M - P - M - M M M - P - P M - - P M - - - - M - - M M - - - -
Aβ-eEPSCs P - M - M - M M - - - M - M - M - - P - P M - - - P - - - P - - - P - - M
Aδ-eEPSCs M - - P M P M - - P M - - - - - - M - - - M M - M - M P P - P - M - P P -
SOM
Gapdh
Cell # 38 39 40 41 42 43 44 45 46 47 48 49 50 51 52 53 54 55 56 57 58 59 60 61 62 63 64 65 66 67 68 69 70 71 72 73
I-EPSCs - - - - - - - - - - - - - - - - - - - - - - - - - - - - - - - - - - - -
I-APs - - - - - - - - - - - - - - - - - - - - - - - - - - - - - - - - - - - -
C-eEPSCs M M M P - - - M - - M P - M - M P - P P P M P - M M - - - - - M P - M -
Aβ-eEPSCs M - M - - - P M - M - - - - P M - - - - M M - P - - M - P - - M - P - M
Aδ-eEPSCs - - M - M P - - P - - - M - - - - M - - P - - - M - - M - P P - - - M -
SOM
Gapdh
Cell # 74 75 76 77 78 79 80 81 82 83 84 85
I-EPSCs - - - - - - - - - - - -
I-APs - - - - - - - - - - - -
C-eEPSCs - M - M - - P M - M - -
Aβ-eEPSCs M P - - - M - - M - M M
Aδ-eEPSCs M - M - M - - - - - - M
SOM
Gapdh
